# *N*^6^-methyladenosine (m^6^A) is an endogenous A3 adenosine receptor ligand

**DOI:** 10.1101/2020.11.21.391136

**Authors:** Akiko Ogawa, Chisae Nagiri, Wataru Shihoya, Asuka Inoue, Kouki Kawakami, Suzune Hiratsuka, Junken Aoki, Yasuhiro Ito, Takeo Suzuki, Tsutomu Suzuki, Toshihiro Inoue, Osamu Nureki, Hidenobu Tanihara, Kazuhito Tomizawa, Fan-Yan Wei

**Author notes:** These authors contributed equally to this study: Chisae Nagiri, Wataru Shihoya, and Asuka Inoue.

## Abstract

About 150 post-transcriptional RNA modifications have been identified in all kingdoms of life. During RNA catabolism, most modified nucleosides are resistant to degradation and are released into the extracellular space. In this study, we explored the physiological role of these extracellular modified nucleosides and found that *N*^6^-methyladenosine (m^6^A), widely known as an epigenetic mark in RNA, acts as a ligand for the adenosine A3 receptor, for which it has greater affinity than unmodified adenosine. Structural modeling defined the amino acids required for specific binding of m^6^A to the A3 receptor. m^6^A is dynamically released in response to cytotoxic stimuli and facilitates type I allergy. Our findings shed light on m^6^A as a signaling molecule with the ability to activate GPCRs, a previously unreported property of RNA modifications.

## INTRODUCTION

Cellular RNA species, including transfer RNA (tRNA), ribosomal RNA (rRNA), and messenger RNA (mRNA), contain a wide variety of post-transcriptional chemical modifications. About 150 species of RNA modifications have been identified in RNA species from all kingdoms of life (Boccaletto et al., 2018). These modifications are indispensable for the localization and stability of RNA molecules, as well as for their physiological function as regulators of translation (Roundtree et al., 2017a; Frye et al., 2016). Dysregulation of RNA modifications is associated with various human disorders, including diabetes, cancer, and mitochondrial diseases (Wei et al., 2011; Wei et al., 2015; Fakruddin et al., 2018; Jonkhout et al., 2017). In contrast to the numerous studies on the biological and pathological functions of RNA modifications, the metabolic fate of modified RNAs is not fully understood.

RNA is constantly undergoing turnover and its degradation is a catabolic process. During RNA breakdown, unmodified nucleosides are recycled for nucleotide biosynthesis through salvage pathways or degraded into uric acid and β-aminosibutyric acid/β-alanine. Alternatively, unmodified nucleosides are released into the extracellular space where they act as signaling molecules. Among unmodified nucleosides, the purine nucleoside adenosine has been intensively studied because of its vital roles in pathophysiological functions. Extracellular adenosine can be generated from hydrolysis of extracellular adenosine triphosphate (ATP) or efflux from the cytosol through equilibrative nucleoside transporters (ENTs) (Eltzschig et al., 2009). The extracellular adenosine binds to four types of purinergic receptors coupled with distinct G proteins, which subsequently regulates second messenger pathways including cAMP and calcium mobilization, and induces a multitude of physiopathological responses including circulation, immune responses, and central nervous systems (Borea et al., 2018).

In contrast to unmodified nucleosides, modified nucleosides are preferentially excreted and cannot enter the salvage pathway due to virtual absence of specific kinases and stable 5’-nucleotide intermediates during enzymatic hydrolysis of oligonucleotides containing modified nucleotides (Uziel et al., 1980; Uziel et al., 1979; Lothrop et al., 1982). Consequently, modified nucleosides are excreted as metabolic end products in urine after circulation in blood (Pane et al., 1992; Willmann et al., 2015; Mandel et al., 1966). To date, some extracellular modified nucleosides have been identified as potential biomarkers of certain diseases, including tumors and AIDS (Borek et al., 1977; Seidel et al., 2006), whether these modified nucleosides can directly activate receptors has remained unexplored.

To test the hypothesis that extracellular modified nucleosides might serve as potential ligands for receptors, we screened a panel of modified nucleosides that were clearly present in human plasma against P1 class purinergic receptors. The screening revealed that *N*^6^-methyladenosine (m^6^A) molecule, one of the most abundant chemical modifications on mRNA, is a selective and potent ligand of the adenosine A3 receptor (A3R). We also identified the downstream signaling pathways mediated by m^6^A and demonstrate that m^6^A binding to the A3R receptor has implications for allergic responses mediated by A3R. Collectively, our findings reveal a previous unknown property of m^6^A as an extracellular signaling molecule and thus provide a starting point for future studies into extracellular modified nucleoside-mediated pathophysiology.

## RESULTS

### Activation of adenosine A3 receptor by extracellular m^6^A

We applied a mass spectrometry-based methodology to quantitatively profile modified nucleosides in human plasma, in which a total of 20 species of modified nucleosides were clearly detected (Fig. S1A-B). When modified nucleosides were classified into subclasses, seven species of modified adenosines, four species of modified uridines, cytidines, and guanosine, and two species modified inosines were detected. Modified nucleosides in these fluids constituted 49% of all nucleosides in human plasma (Fig. 1A). These modified nucleosides were also abundant in serum, aqueous humor, vitreous humor and urine, not only in human but also in non-human mammals such as mouse and rabbit (Fig. 1A, S1C).

**Figure 1.**
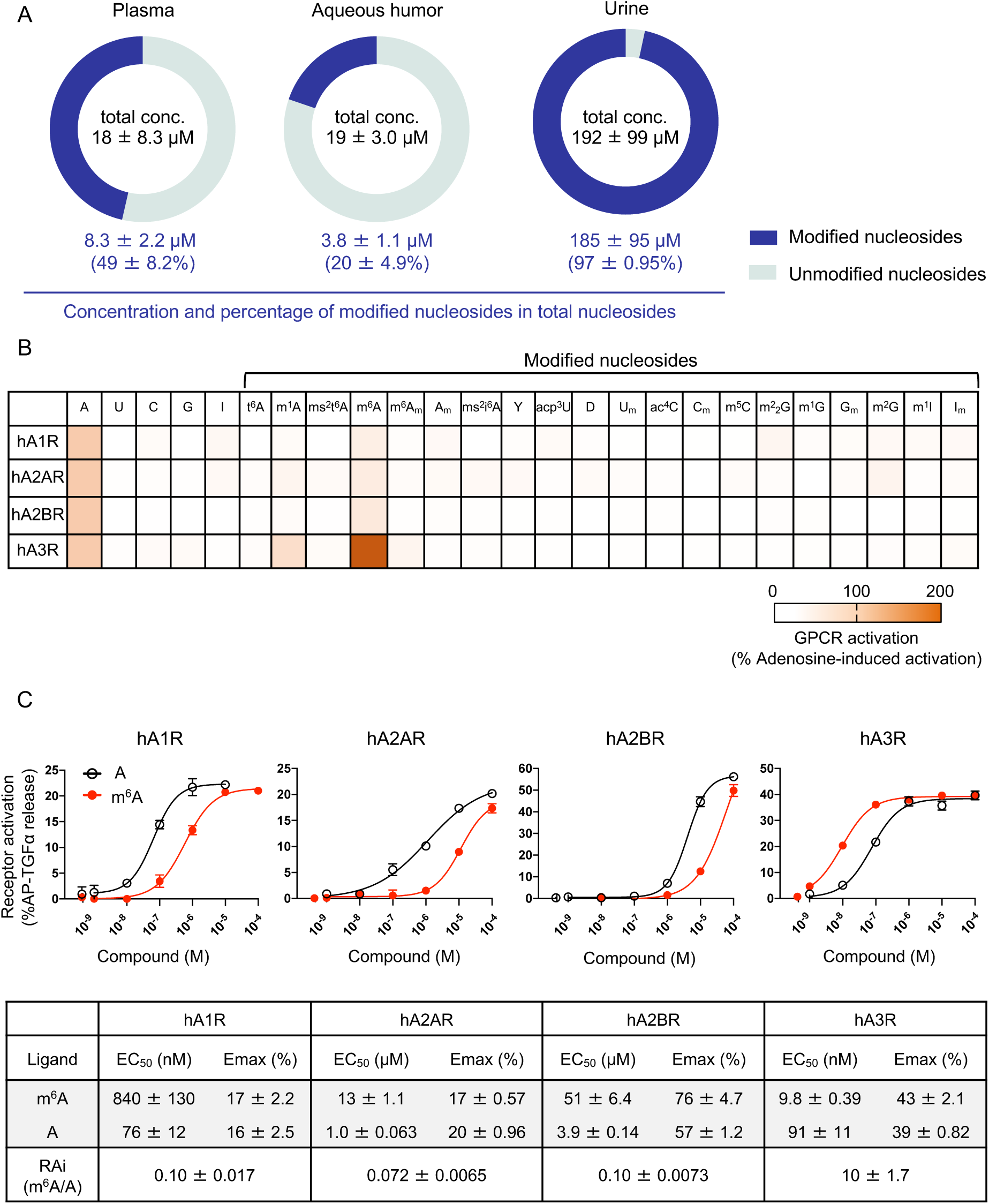
Activation of A3R by extracellular m^6^A. (A) Substantial levels of modified nucleosides were detected in human extracellular fluids. n = 7 for plasma, n = 9 for aqueous humor, and n = 12 for urine. Total nucleoside concentration in each extracellular fluid is shown in the middle of each pie chart, and the concentration and amount of modified nucleosides as a percentage of total nucleosides is shown below each pie chart. Data are shown as means ± SD. (B) Comparisons of adenosine receptor activation by different nucleoside derivatives in human plasma, based on AP-TGFα release percentages. Values were obtained by TGFα-shedding assay using test compounds at 100 nM for A1R and A3R, 1 µM for A2AR, and 10 µM for A2BR. m^6^A had the most pronounced effect on A3R. Values are shown as an average of 3-10 independent experiments. (C) TGFα-shedding response curves of m^6^A and adenosine for each adenosine receptor subtype. m^6^A exhibited higher A3R activation capacity than adenosine. Symbols and error bars are means ± SEM of representative experiments with each performed in triplicate. Parameters (means ± SEM) obtained from three independent experiments are shown at the right panel. See also Figure S1.

The wide variety of extracellular modified nucleosides prompted us to speculate that these molecules might act as ligands for receptors. Among nucleoside receptors, adenosine and its derivatives have been extensively studied as natural ligands of a subset of G-protein-coupled receptors (GPCRs) known as P1 class purinergic adenosine receptors (Jacobson et al., 2006). Adenosine receptors (ARs) have been grouped into four subtypes (A1, A2A, A2B, and A3), all of which are activated by extracellular adenosine. Using a TGF-α shedding assay (Inoue et al., 2012), we screened 20 modified nucleosides to determine if they would activate ARs. We found that m^1^A, m^6^A, and m^6^Am were capable of activating ARs, with m^6^A having the highest activation level for A3R (Fig 1B). A detailed pharmacological analysis revealed that the potency of m^6^A for A3R activation was higher than that of adenosine (EC_50_, m^6^A: 9.8 ± 0.39 nM; adenosine: 91 ± 11 nM: Fig 1C). The potency of m^6^A as an activator of A3R was comparable to those of well-known synthetic A3R agonists (Fig. S1D). To quantitatively analyze the potency of m^6^A relative to adenosine, we calculated relative intrinsic activity (RAi) values by dividing the E_max_/EC_50_ value of m^6^A by that of adenosine (Inoue et al., 2019; Ehlert et al., 1999). The RAi value of m^6^A vs. adenosine was 10 ± 1.7 for A3R but much lower for adenosine receptors (A1R: 0.10 ± 0.017, A2AR: 0.072 ± 0.0065, A2BR: 0.10 ± 0.0073; Fig. 1). m^6^A-induced A3R activation was confirmed by two other GPCR assays that detect proximal signaling events including G_i1_ dissociation (Fig. S1E) and β-arrestin recruitment (Inoue et al., 2019) (Fig. S1F). In both experiments, the potency of m^6^A was markedly higher than that of adenosine. These findings show that m^6^A is a potent and selective ligand for A3R; moreover, our results show that m^6^A, along with other modified nucleosides, constitutes a previously uncharacterized category of metabolites.

### The origin of extracellular m^6^A

m^6^A is one of the most abundant modifications in mRNA, but it is also present in other RNA species including rRNA, U6 small nuclear RNA (snRNA), and long non-coding RNA (lncRNA) (Roundtree et al., 2017a; Gilbert et al., 2016; Zhao et al., 2017). All methyl groups in the methylated purine and pyrimidine moieties of RNA molecules originate from direct transfer of the methyl group of methionine (Mandel et al., 1963). To determine the origin of extracellular m^6^A, we utilized stable isotope-labeled methionine (^13^C-Met), which can be incorporated into RNA as a methyl donor. Cells were treated with ^13^C-Met for 12 h, followed by a chase period of 48 h (Fig. 2A). A decrease in the level of ^13^C-m^6^A in RNA was clearly associated with an increase in the extracellular level of ^13^C-m^6^A, suggesting that extracellular ^13^C-m^6^A was derived from RNA as a result of RNA turnover (Fig. 2B). We also injected ^13^C-Met into the anterior chamber of rabbit eyes (Fig. S2A) and detected ^13^C-m^6^A in eye fluids as an extracellular metabolite (Fig. S2B), as well as in RNA from ocular tissues (Fig. S2C).

**Figure 2.**
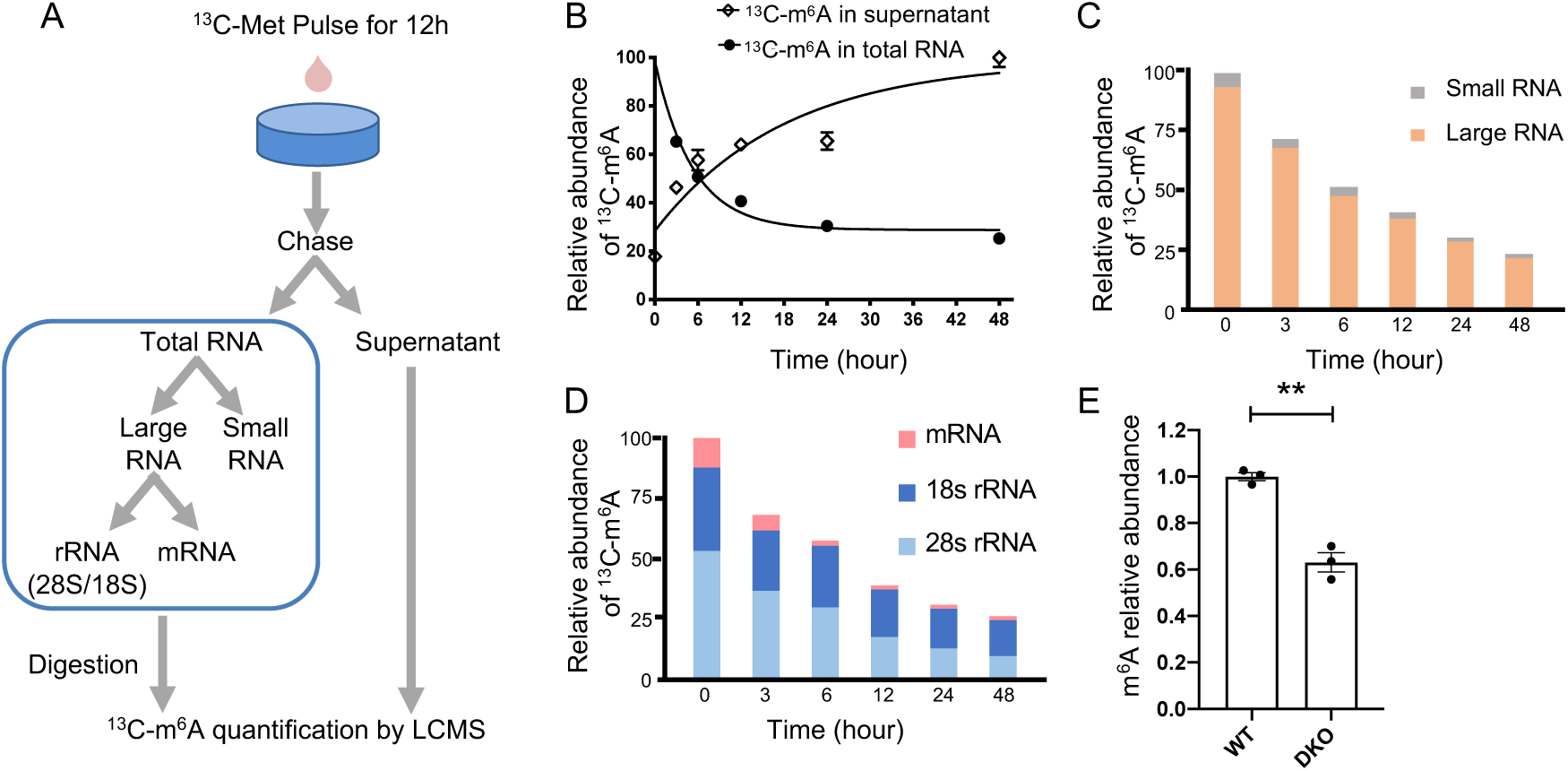
Extracellular m^6^A is derived from multiple RNA species. (A) Schematic of the pulse-chase labeling of m^6^A with stable isotope labeled methionine. (B) Relative abundance of ^13^C-labeled m^6^A in total RNA and supernatant (sup) in cells treated with stable isotope (^13^C)-labeled methionine. ^13^C-m^6^A in total RNA exhibited one-phase decay, with a gradual increase in the extracellular ^13^C-m^6^A level. Symbols and error bars represent means and SEM, respectively, of two independent experiments with each performed in triplicate. (C) Relative abundance of labeled m^6^A in large RNA (> 200 nt) and small RNA (< 200 nt), as determined by pulse-chase labeling. More than 90% of m^6^A was present in large RNA. (D) Large RNA was further fractioned into mRNA, 28S, and 18S rRNA, and the decay of ^13^C-m^6^A in these RNA species was examined. At the beginning of the chase period, rRNA and mRNA contained 87% and 13% of labeled m^6^A, respectively. Half-lives were 5.5 and 4.9 h for 18S and 28S rRNA, respectively, vs. 2.2 h for mRNA. (E) Extracellular m^6^A level at steady state decreased to 63% in cells lacking both *ZCCHC4* and *METTL5* (DKO), which are responsible for m^6^A modifications of 28S and 18S rRNA, respectively. **P < 0.01 vs. control (unpaired two-tailed Student’s *t*-test). See also Figure S2.

Next, we measured the relative abundance of ^13^C-m^6^A in major RNA species and the kinetics of its decline. ^13^C-m^6^A containing RNA was separated into < 200 nt small RNA (tRNA and snRNA) and > 200 nt large RNA (rRNA and mRNA), which was further separated into 18S rRNA, 28S rRNA, and mRNA (Fig. 2A). At the beginning of the chase period, more than 90% of ^13^C-m^6^A in total RNA was found in large RNAs (Fig. 2C), of which 87% was present in rRNA and 13% in mRNA (Fig. 2D), indicating that most m^6^A had been incorporated into rRNAs. During the chase period, the levels of ^13^C-m^6^A in both 18S rRNA and 28S rRNA gradually declined, with half-lives of 5.5 and 4.9 h, respectively. By contrast, turnover of ^13^C-m^6^A in mRNA was very fast, with a half-life of 2.2 h. Therefore, it is likely that extracellular m^6^A is derived from multiple RNA species including mRNA and rRNA. In line with this hypothesis, the extracellular level of m^6^A significantly decreased to 63% in cells lacking both ZCCHC4 and METTL5 (Fig. 2E), which are responsible for m^6^A modification of 28S and 18S rRNA, respectively (Fig. S2D-F) (Ma et al., 2019; van Tran et al., 2019).

### Stimulation-dependent release of m^6^A

The extracellular adenosine level changes dynamically in response to various stresses, e.g., cytotoxic stimulation and hypoxia (Jackson et al., 2017; Saito et al., 1999). To determine whether extracellular m^6^A is also affected by external stimuli, we challenged cells with various cytotoxic reagents and measured the amount of extracellular m^6^A by mass spectrometry. Mitomycin C (MMC), staurosporine, and hydrogen peroxide (H_2_O_2_) all increased extracellular m^6^A in a dose-dependent manner (Fig. 3A-D), and the H_2_O_2_-induced increase was decreased to 63% in cells lacking the two enzymes required for m^6^A modification of rRNA (Fig. 3E). Notably, the extracellular levels of other modified adenosines did not increase, but rather decreased, when cells were treated with cytotoxic reagents (Fig. 3A). Furthermore, extracellular m^6^A and adenosine exhibited distinct release patterns: extracellular adenosine levels rapidly increased but returned to baseline 24 h after H_2_O_2_ treatment, whereas extracellular m^6^A levels gradually increased and peaked 24 h after H_2_O_2_ treatment (Fig. 3F). In addition to cytotoxic conditions, we also subjected cells to hypoxic conditions and monitored the extracellular m^6^A level. Hypoxia markedly decreased the extracellular levels of m^6^A and other modified adenosines (Fig. S3A-B). The stimulation-dependent release of m^6^A in culture cells prompted us to examine m^6^A release *in vivo*. Specifically, we monitored m^6^A and adenosine in serum and tissue interstitial fluids in a mouse model of lipopolysaccharide (LPS)-induced shock. After LPS injection, the m^6^A level increased in serum, liver, spleen, and lung (Fig. 3G), whereas the adenosine level did not change significantly, except in liver (Fig. 3H). These results suggest that m^6^A is actively released in response to cytotoxic stimulation *in vivo*, and that its pathophysiological role in the stress response is distinct from that of adenosine.

**Figure 3.**
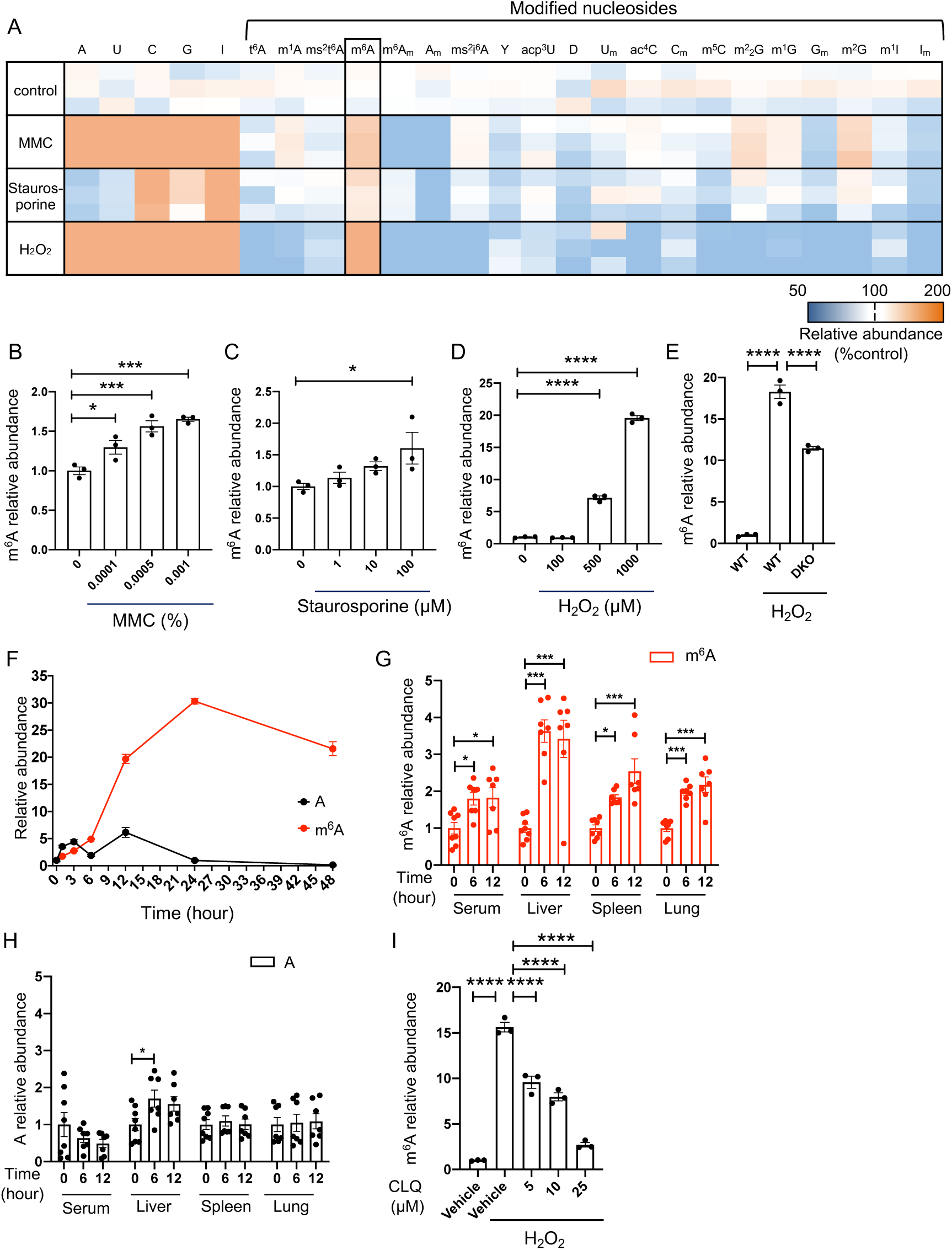
m^6^A is dynamically released upon cytotoxic stimulation both *in vitro* and *in vivo*. (A) Heatmap of the differentially released extracellular nucleoside derivatives in HEK 293 supernatants following various cytotoxic treatments: mitomycin C (MMC, 0.001%), staurosporine (0.1 µM), and hydrogen peroxide (H_2_O_2_, 1 mM). Three (n = 3) independent experiments were performed. Representative results are shown. (B-D) m^6^A was released into supernatants of HEK 293 cells 24 h after cytotoxic stimulation with (B) MMC, (C) staurosporine, and (D) H_2_O_2_, in a dose-dependent manner. *P < 0.05, ***P < 0.001, and ****P < 0.0001 vs. non-treated group (one-way ANOVA followed by a Dunnett’s multiple comparison test). Bar graph representative of two independent experiments with each experiment containing three biological replicates. Data are means ± SEM. (E) Extracellular m^6^A surge after exposure to 1 mM H_2_O_2_ decreased to 63% in cells lacking both ZCCHC4 and METTL5 (DKO) cells. ****P < 0.0001 vs. 1 mM H_2_O_2_ (one-way ANOVA followed by a Dunnett’s multiple comparison test). Bar graph representative of two (n = 3) independent experiments. Data are means ± SEM. (F) Extracellular m^6^A and adenosine exhibited distinct release patterns upon H_2_O_2_ treatment. Symbols and error bars represent means and SEM, respectively, of two independent experiments with each performed in triplicate. (G and H) Abundance of (G) m^6^A and (H) adenosine in serum and multiple tissue interstitial fluids in an LPS-induced shock mouse model. After LPS stimulation, the m^6^A level was increased in all extracellular fluids, whereas adenosine only exhibited a slight increase in liver interstitial fluids. (I) Cytotoxicity-induced release of m^6^A was dose-dependently inhibited by pretreatment with chloroquine (CLQ), a lysosome inhibitor. The indicated concentrations of CLQ were added 12 h before 24 h H_2_O_2_ treatment. ****P < 0.0001 vs. 1 mM H_2_O_2_ group (one-way ANOVA followed by a Dunnett’s multiple comparison test). Bar graph is representative of three (n = 3) independent experiments. Data are means ± SEM. See also Figure S3.

Next, we sought to determine the factors that are responsible for the generation of extracellular m^6^A. The major RNA degradation pathways in the cell are Xrn1/2-mediated 5’-3’ RNA degradation and exosome-mediated 3’-5’ RNA degradation (Houseley et al., 2009). Knockdown of key ribonucleases in the two pathways revealed that 5’-3’ exoribonuclease 1 (XRN1) is involved in the generation of extracellular m^6^A. At steady state, the extracellular m^6^A level decreased by 45 % upon silencing of XRN1 (Fig. S3C). Silencing of XRN1 also significantly decreased the level of extracellular m^6^A upon H_2_O_2_ stimulation, although the reduction was moderate (19% reduction) (Fig. S3D), suggesting that an XRN1-independent RNA degradation pathway is also involved in the generation of extracellular m^6^A.

Besides ribonuclease-mediated degradation, the lysosome is the major organelle for RNA degradation (Arsenis et al., 1970; Fujiwara et al., 2013). To determine whether the lysosome is involved in the generation of extracellular m^6^A, we treated cells with the lysosome inhibitor chloroquine and exposed them to H_2_O_2_. As with XRN1 knockdown, chloroquine treatment decreased the level of extracellular m^6^A (Fig. S3E). Notably, chloroquine dramatically decreased m^6^A release induced by H_2_O_2_ treatment in a dose-dependent manner (Fig. 3I). The same drastic decrease was observed upon treatment with another lysosomal inhibitor, concanamycin A (Fig. S3F). By contrast, inhibition of lysosomal protease activity with E64d had no effect on extracellular m^6^A release (Fig. S3G). Collectively, our results demonstrated that basal m^6^A release is mediated by both XRN1 and lysosomes, whereas stimulation-dependent m^6^A release is mainly regulated by lysosomes.

### m^6^A-mediated signaling transduction and its implications for the allergic response

To determine whether extracellular m^6^A is capable of activating intracellular signaling transduction via A3R, we expressed A3R in HEK293 cells and stimulated them with m^6^A. Application of m^6^A induced a rapid phosphorylation of extracellular signal-regulated kinase 1/2 (ERK1/2), which was abolished by the A3R antagonist or the G_i_ inhibitor pertussis toxin (PTX) (Fig. S4A). Calcium is a key second messenger downstream of A3R activation (Shneyvays et al., 2004). Application of m^6^A rapidly induced an intracellular calcium transient that was effectively abolished by the A3R antagonist (Fig. S4B).

Next, we sought to reveal the physiological role of m^6^A as a ligand of A3R. A3R is broadly expressed in tissues, with relatively high expression in immune cells. In the immune system, A3R activation contributes to the type I allergic response by facilitating mast cell degranulation (Reeves et al., 1997; Borea et al., 2015). To determine whether m^6^A is capable of activating mast cells, we treated rat mast cell-derived RBL-2H3 cells with m^6^A in the presence of antigen (IgG). m^6^A potentiated mast cell degranulation in a dose-dependent manner (Fig. 4A). Mechanistically, m^6^A evoked typical G_i_-mediated signaling, including suppression of cAMP production and resulted in an intracellular calcium transient through A3R (Fig. 4B-C). Furthermore, to confirm that m^6^A can facilitate the type I allergy response *in vivo*, we used a passive cutaneous anaphylaxis (PCA) model (Ovary et al., 1958). Application of m^6^A *in vivo* significantly enhanced the allergic reaction in a dose-dependent manner (Fig. 4D-E), and this effect was potently inhibited by treatment with A3R antagonist (Fig. 4F).

**Figure 4.**
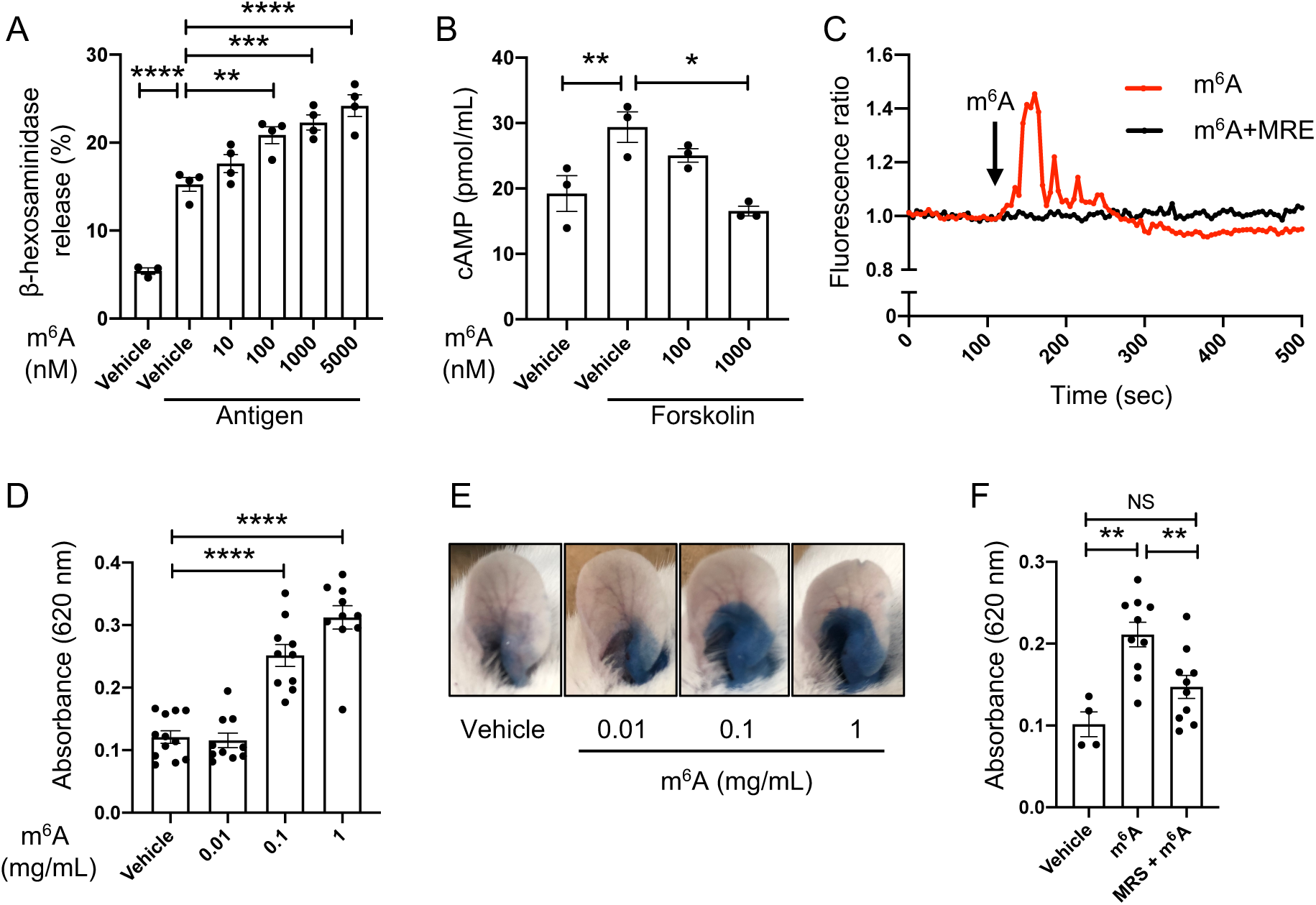
m^6^A facilitates a type I allergy response. (A) m^6^A increased antigen-mediated mast cell degranulation in a dose-dependent manner. **P < 0.01, ***P < 0.001 and ****P < 0.0001 vs. antigen without m^6^A (one-way ANOVA followed by a Dunnett’s multiple comparison test). Bar graph is representative of the results of three (n = 4) independent assays of the *in vitro* degranulation assay using RBL-2H3 cells. Data are means ± SEM. (B) m^6^A suppressed forskolin-mediated cAMP production in RBL-2H3 cells. *P < 0.05 and **P < 0.01 vs. forskolin without m^6^A group (one-way ANOVA followed by a Dunnett’s multiple comparison test). Bar graph is representative of two (n = 3) independent experiments. Data are means ± SEM. (C) Intracellular calcium transient in RBL-2H3 cells in response to 5 µM m^6^A. m^6^A-induced calcium transients were effectively blocked by pretreatment with the A3R antagonist (1 μM MRE 3008F20; MRE). The transients of ten representative cells are shown. (D) m^6^A promotes an antigen-induced type I allergy response in a murine PCA model *in vivo*. The effect is dose-dependent. Absorbance of extravasated dye is shown. ****P < 0.0001 vs. antigen without m^6^A group (one-way ANOVA followed by a Dunnett’s multiple comparison test; n = 10). Data are means ± SEM. (E) Representative photos of ears showing dye extravasation. (F) m^6^A-mediated facilitation of the allergic response was blocked by A3R antagonist MRS 1191 (MRS). **P < 0.01 (one-way ANOVA followed by a Tukey’s multiple comparisons test; n = 4-10). Data are means ± SEM. See also Figure S4.

### Molecular basis of selective A3R activation by m^6^A

Finally, we sought to elucidate the molecular basis underlying m^6^A-mediated activation of A3R. The structures of the orthosteric pockets in A1R and A2AR are very similar (Draper-Joyce et al., 2018; Carpenter et al., 2016), suggesting that this feature is highly similar among adenosine receptors. Among the four adenosine receptor subtypes, A3R and A1R share 41% identity at the amino acid sequence level, and both receptors bind to G_i/o_ proteins (Fig. S5A). Given the similarities between A3R and A1R, we performed homology modeling of A3R based on the A1R structure (Fig. 5A upper), which was solved by cryo-electron microscopy in complex with adenosine and G_i_ protein (PDB ID 6D9H) (Draper-Joyce et al., 2018). The A3R homology model indicated that the orthosteric binding pocket for m^6^A is highly conserved between A1R, A2AR, and A3R, except for the regions in between extracellular loop 2 (ECL2) and transmembrane (TM) domains 6 and 7 (Fig. 5B, Fig. S5A). Notably, A3R contains hydrophobic V169^5.30^ and I253^6.58^ at ECL2 and TM6, whereas the corresponding amino acids in A1R and A2AR are hydrophilic glutamic acid and threonine (E172^5.30^ and T257^6.58^ for A1R; E169^5.30^ and T256^6.58^ for A2AR). Moreover, a recent study showed that A1R T270^7.35^ and A2AR M270^7.35^, which correspond to L264^7.35^ at TM7 in A3R, confer ligand selectivity (Cheng et al., 2017). The A3R homology model demonstrated that the methyl group of m^6^A can form a close van der Waals interaction with the hydrophobic residues V169^5.30^, I253^6.58^, and L264^7.35^ of A3R (Fig. 5A, lower). By contrast, adenosine is relatively distant from these three residues due to its lack of a methyl group (Fig. 5A, lower). These observations showed that an addition of a methyl group at the nitrogen-6 position of adenosine stabilizes the hydrophobic region of A3R, and that this contributes to the receptor’s selectivity for m^6^A.

**Figure 5.**
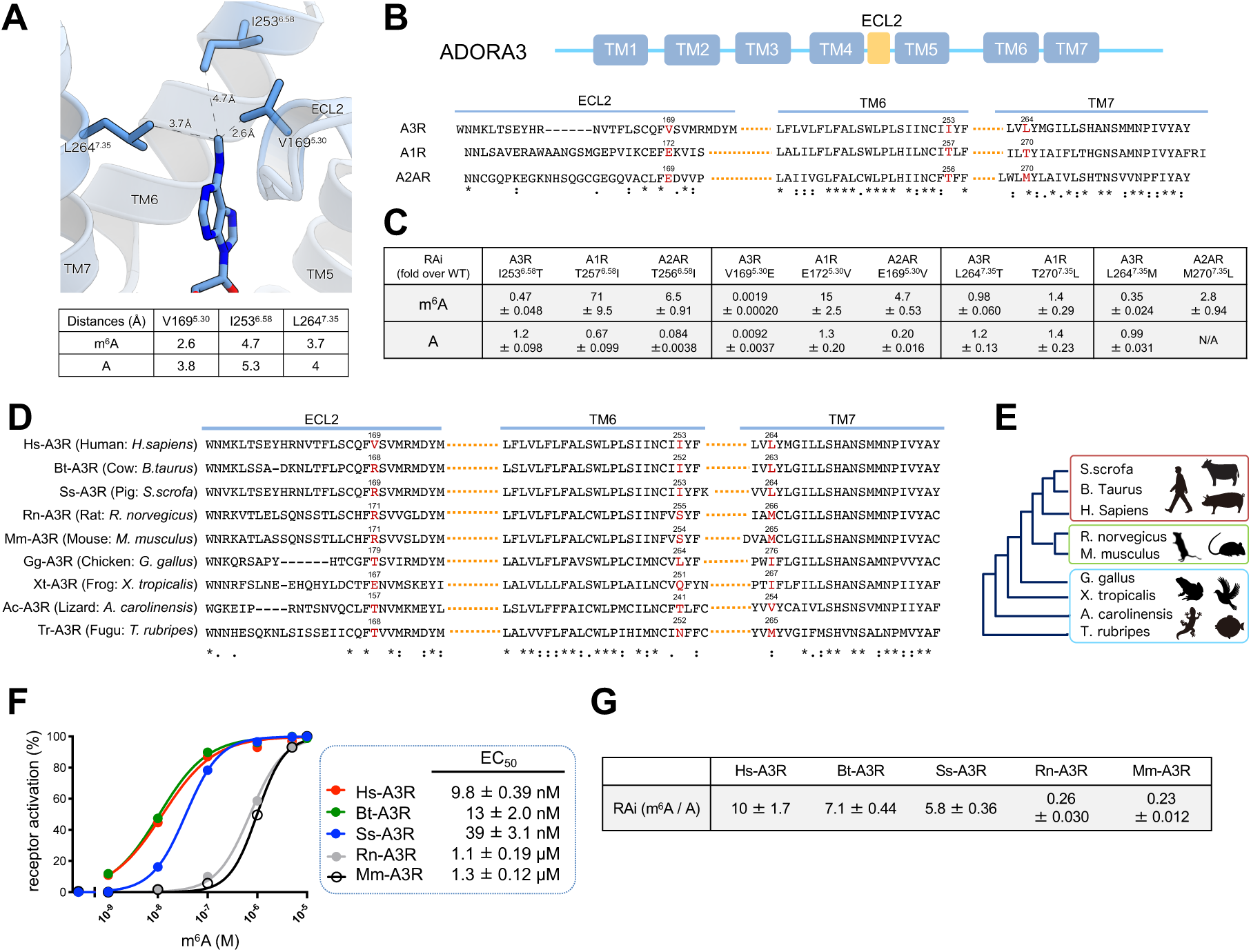
Molecular basis of selective A3R activation by m^6^A. (A) Homology-modeled structure of the m^6^A binding site in the human adenosine A3 receptor (PDB code: 6D9H, upper panel), and estimated distances of m^6^A and adenosine from V169^5.30^, I253^6.58^ and L264^7.35^ in A3R (lower panel). (B) Amino acid alignment of the orthosteric binding pocket for m^6^A in A1R, A2AR, and A3R, showing high conservation except for the regions in between extracellular loop 2 (ECL2), transmembrane (TM) domains 6 and 7. (C) Relative RAi values of m^6^A and adenosine for A3R, A1R, and A2AR mutants, as determined by the TGF-α shedding assay. RAi values are expressed as fold change of the values for WT. I253^6.58^T and L264^7.35^M mutations selectively reduced A3R activation by m^6^A, and the corresponding A1R T257^6.58^I, A2AR T256^6.58^I, and A2AR M270^7.35^L mutants showed an increase in the receptor sensitivity for m^6^A. Data are expressed as means ± SEM of three independent experiments with each performed in triplicate. Values used for calculating RAi are shown in Figure S5B. (D) Alignment of the ECL2, TM6, and TM7 of human (Hs), cow (Bt), pig (Ss), rat (Rn), mouse (Mm), chicken (Gg), frog (Xt), lizard (Ac), and *Fugu* (Tr), highlighting evolutionary variability of the three residues. (E) Phylogenetic tree of A3R across vertebrates obtained by amino acid sequence alignment using UniProt. (F) Concentration-response curves of m^6^A for mammalian A3R, as determined by the TGF-α shedding assay. The original values were fitted to an operational model to obtain EC_50_ values. Symbols and error bars are means ± SEM of representative experiments with each performed in triplicate. Parameters of the EC_50_ values (means ± SEM) obtained from three independent experiment are shown to the right. The %AP-TGFα release of each A3R at 10^-5^ M was defined as 100%. EC_50_ values differed largely between large mammals (human, cow, and pig) and small mammals (rat and mouse). (G) Relative RAi values of m^6^A vs. adenosine for A3R in the indicated mammals. A3R activation capacity of m^6^A was higher than that of adenosine in large mammals but lower than that of adenosine in small mammals. Data are expressed as mean ± SEM of three independent experiments with each performed in triplicate. See also Figure S5.

To experimentally validate the result of homology modeling, we mutated these residues in WT A3R (V169^5.30^, I253^6.58^, and L264^7.35^) to the corresponding residues in A1R or A2AR, and tested whether the mutations could selectively decrease m^6^A-mediated A3R activation (Fig. 5C, Fig. S5B). Among the series of mutant A3Rs, the I253^6.58^T and L264^7.35^M mutations selectively reduced A3R activation by m^6^A (RAi = 0.47 ± 0.048 for I253^6.58^T; RAi = 0.35 ± 0.024 for L264^7.35^M), but not by adenosine (RAi = 1.2 ± 0.098 for I253^6.58^T; RAi = 0.99 ± 0.031 for L264^7.35^M). On the other hand, the V169^5.30^E mutation abolished both m^6^A- and adenosine-mediated A3R activation (RAi = 0.0019 ± 0.00020 for m^6^A; RAi = 0.0092 ± 0.0037 for adenosine). In addition, we mutated the residues in A1R or A2AR to the corresponding residues in A3R, and tested whether the mutations could increase the response of A1R and A2AR to m^6^A. Indeed, the A1R T257^6.58^I and A2AR T256^6.58^I mutants exhibited an increase in receptor sensitivity toward m^6^A (RAi = 71 ± 9.5 and 6.5 ± 0.91, respectively) and a decrease in sensitivity toward adenosine (RAi = 0.67 ± 0.099 and 0.084 ± 0.0038, respectively). Likewise, the A2AR M270^7.35^L mutant also exhibited an increase in receptor sensitivity for m^6^A (RAi = 2.8 ± 0.94). Moreover, the A2AR E169^5.30^V mutant showed an increase in receptor sensitivity to m^6^A (RAi = 4.7 ± 0.53) and a concomitant decrease in receptor sensitivity to adenosine (RAi = 0.20 ± 0.016), whereas the A1R E172^5.30^V mutant exhibited an increase in receptor sensitivity to both m^6^A and adenosine (RAi = 15 ± 2.5 and 1.3 ± 0.20, respectively). These results suggest that A3R I253^6.58^ and L264^7.35^ are required for the selectivity toward m^6^A, whereas A3R V169^5.30^ is required for the affinity for m^6^A.

The important role of I253^6.58^, L264^7.35^, and V169^5.30^ in m^6^A-mediated A3R activation prompted us to investigate whether these residues are evolutionarily conserved. Alignment of A3R across vertebrates revealed that these three residues changed over the course of evolution, whereas the other residues constituting the orthosteric site are highly conserved (Fig. 5D-E, Fig. S5C). I253 and L264 are only conserved in human, cow, and pig; by contrast, mouse and rat have serine and methionine at the corresponding residues. V169 is only present in human A3R (hs-A3R), whereas other mammals have arginine at the corresponding residue (Fig. 5D). Moreover, the variability of these residues is even higher among non-mammalian vertebrates than among mammals. Consistent with this, a GPCR assay revealed that the potency of m^6^A-mediated A3R activation gradually increased with evolutionary closeness to human (EC_50_: not applicable in non-mammalian vertebrates; 1.3 ± 0.12 µM for *M. musculus* A3R (Mm-A3R); 1.1 ± 0.19 µM for *R. norvegicus* A3R (Rn-A3R); 39 ± 3.1 nM for *S. scrofa* (Ss-A3R); 13 ± 2.0 nM for *B. taurus* A3R (Bt-A3R); and 9.8 ± 0.39 nM for *H. sapiens* (Hs-A3R); Fig. 5F). Notably, the selectivity for m^6^A vs. adenosine also exhibited the same evolutionary pattern (RAi = 0.23 ± 0.012 for Mm-A3R; 0.26 ± 0.030 for Rn-A3R; 5.8 ± 0.36 for Ss-A3R; 7.1 ± 0.44 for Bt-A3R; and 10 ± 1.7 for Hs-A3R, Fig. 5G, Fig. S5D). Our data suggest that the acquisition of m^6^A reactivity emerged as a consequence of A3R evolutionary changes in particular A3R residues.

## DISCUSSION

To date, over 150 species of RNA modifications have been identified in all domains of life. It has been known for decades that these modified nucleosides are excreted as metabolic end products of RNA breakdown (Pane et al., 1992; Willmann et al., 2015; Mandel et al., 1966). However, most studies have focused on their potential clinical applications as biomarkers (Borek et al., 1977; Seidel et al., 2006). In the present study, we systematically explored the potential role of extracellular modified nucleosides as signaling molecule. As a proof-of-concept, we screened various modified nucleosides against P1 purinergic receptors, and found that some modified nucleosides acted as ligands for ARs. m^6^A was the most selective and potent ligand for A3R even compared to unmodified adenosine. These findings demonstrate that modified nucleosides are not just byproducts of RNA degradation released into the extracellular space, but can act as signaling molecules that regulate physiological functions via their binding to specific receptors on the cell membrane.

RNA modifications have recently emerged as a new research field known as epitranscriptomics. Among these modifications, m^6^A is one of the most abundant modifications of mRNA, and serves as an epigenetic landmark for post-transcriptional gene regulation including alternative splicing, export, structure, stability, processing, metabolism, and translation (Xiao et al., 2016; Roundtree et al., 2017b; Liu et al., 2017; Wang et al., 2014; De Jesus et al., 2019; Wang et al., 2015). Consequently, the m^6^A modification is involved in multiple biological functions including immune response and learning and memory, as well as in tumorigenesis (Winkler et al., 2019; Shi et al., 2018; Lan et al., 2019). In addition to the regulatory roles of the m^6^A modification in RNA function, our results demonstrate that extracellular m^6^A is generated from RNA catabolism and activates A3R as signaling molecule. Furthermore, our homology modeling revealed that the selectivity of m^6^A toward A3R is mediated by specific hydrophobic amino acids located in the substrate binding pocket of A3R. In line with this observation, substitution of these hydrophobic amino acids with neutral or hydrophilic amino acids reduced A3R selectivity for m^6^A. Intriguingly, these residues show species variability in vertebrates. The potency of m^6^A and the selectivity of m^6^A vs. adenosine for A3R activation gradually increased with evolutionarily closeness to humans and with changes in amino acids in the substrate binding pocket of A3R of those of human. Therefore, it is probable that the selectivity toward m^6^A was acquired over the course of evolution as the consequence of changes in key A3R residues.

The release of unmodified and modified nucleosides into the extracellular space is highly dynamic after cells are exposed to external stress such as H_2_O_2_, but the pathways leading to release appear to be different for unmodified and modified nucleosides. It is well-established that extracellular adenosine stems from hydrolysis of ATP, particularly under cytotoxic conditions (Jackson et al., 2017). By contrast, the release of m^6^A appears to involve lysosome-dependent degradation. Because lysosome contains abundant acid nuclease and phosphatase, it is likely that most of m^6^A is generated in lysosome through degradation of macro RNAs. It remains unclear why release of m^6^A but not that of other modified nucleosides is selectively increased by external stress. Further study is needed to identify the lysosomal enzymes that are responsible for the generation and release of m^6^A.

It is worthwhile noting that *N*^6^-methyl-AMP (*N*^6^-mAMP), which is also possibly generated by RNA catabolism, has been detected in plant cells and human cancer cells (Chen et al., 2018). Although the amount of intracellular *N*^6^-mAMP is far lower than that of AMP (Chen et al., 2018), it is conceivable that some *N*^6^-mAMP is released from cells and hydrolyzed to m^6^A upon cytotoxic stimulation.

Although most RNA modifications are catalyzed by corresponding modifier enzymes, RNA modifications can be formed by spontaneous chemical reactions resulting from RNA oxidative damage. For example, 8-hydroxyguanosine (8-OHG) is a major product of RNA oxidation (Fiala et al., 1989; Nunomura et al., 1999) and is detected in human extracellular fluid, including serum and cerebrospinal fluid, where its presence is likely associated with cellular oxidative stress (Gmitterová et al., 2018). A previous study reported that 8-OHG might act as a modulator for toll-like receptors (TLRs), but only in the presence of oligoribonucleotides (Shibata et al., 2016). Given the selective and potent activition of A3R by m^6^A and its implication in mast cells, it is likely that m^6^A and other modified nucleosides might have synergistic effects on the immune response.

A limitation of the current work is that it is technically difficult to manipulate the amount of extracellular m^6^A especially *in vivo*, because genetic deletion of m^6^A-modifying enzymes would impact the function of RNA itself. Development of novel tools, such as neutralizing antibodies against extracellular m^6^A, might help to clarify the physiological functions of m^6^A. In addition, because murine A3R is only moderately activated by m^6^A, evaluating the role of m^6^A in wild-type mice might underestimate its function in human. It will be necessary to generate a humanized A3R mouse model to understand fully the physiological role of m^6^A. In summary, our study presents biochemical and structural evidence that extracellular m^6^A is generated from RNA catabolism and is a potent and selective ligand of A3R. Our study provides a framework for understanding the roles played by the complex world of extracellular modified nucleosides.

## ACKNOWLEDGEMENTS

We are grateful for suggestions and technical supports from T. Chujo and all other members of the Department of Molecular Physiology, Kumamoto University. We thank N. Maeda, H. Masumoto and Y. Takahata for technical assistance and Y. Tashiro, A. Yoshinaga and T. Matsunaga for sample preparation. This work was supported by JSPS KAKENHI grant 18H02599 (F.-Y. W.); 18K19521 (F.-Y. W.); 20H05309 (F.-Y. W.); 18H02865 (K. T.); 17905074 (K. T.); 18959602 (K. T.); 20K18371 (A. O.); 17K08264 (A. I.); and by grants from the Japan Science and Technology Agency (JST), SAKIGAKE JPMJPR1532 (F.-Y. W.); the Japan Agency for Medical Research and Development (AMED), 17935694 (K. T.); the PRIME JP18gm5910013 (A. I.); and the LEAP JP18gm0010004 (A. I. and J. A.); the Takeda Science Foundation (K. T. and F.-Y. W.); and the Uehara Memorial Foundation (F.-Y. W.); the start-up research funding of TUMUG support project conducted by Tohoku University Center for Gender Equality Promotion (A. O.); and the Japan Medical Women’s Association Foundation (A. O.)

## AUTHOR CONTRIBUTIONS

A. O. and F.-Y. W. contributed to the overall conceptualization, design, and management of the study and analyzed the data. C. N. and W. S. performed and analyzed modelling of the m^6^A bound to the A3 receptor. A. I. performed GPC used in this study. Y. I., T. I., and H. T. prepared samples. K. T., N. O. and J. A. supervised the study. A. O., C. N., W. S., A. I., and F.-Y. W. wrote the paper.

## DECLARATION OF INTERESTS

The authors declare no competing interests.

## METHODS

### KEY RESOURCES TABLE

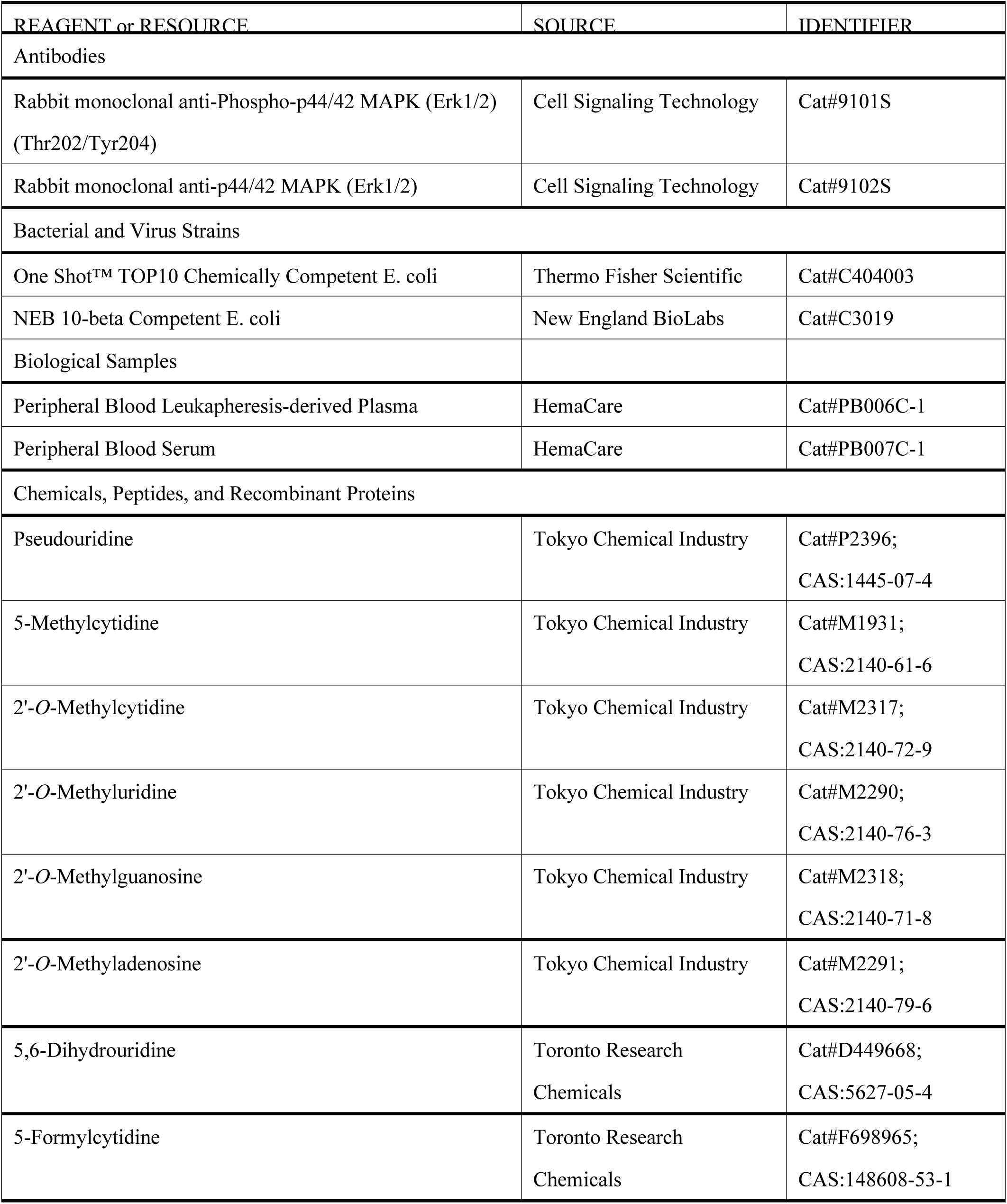

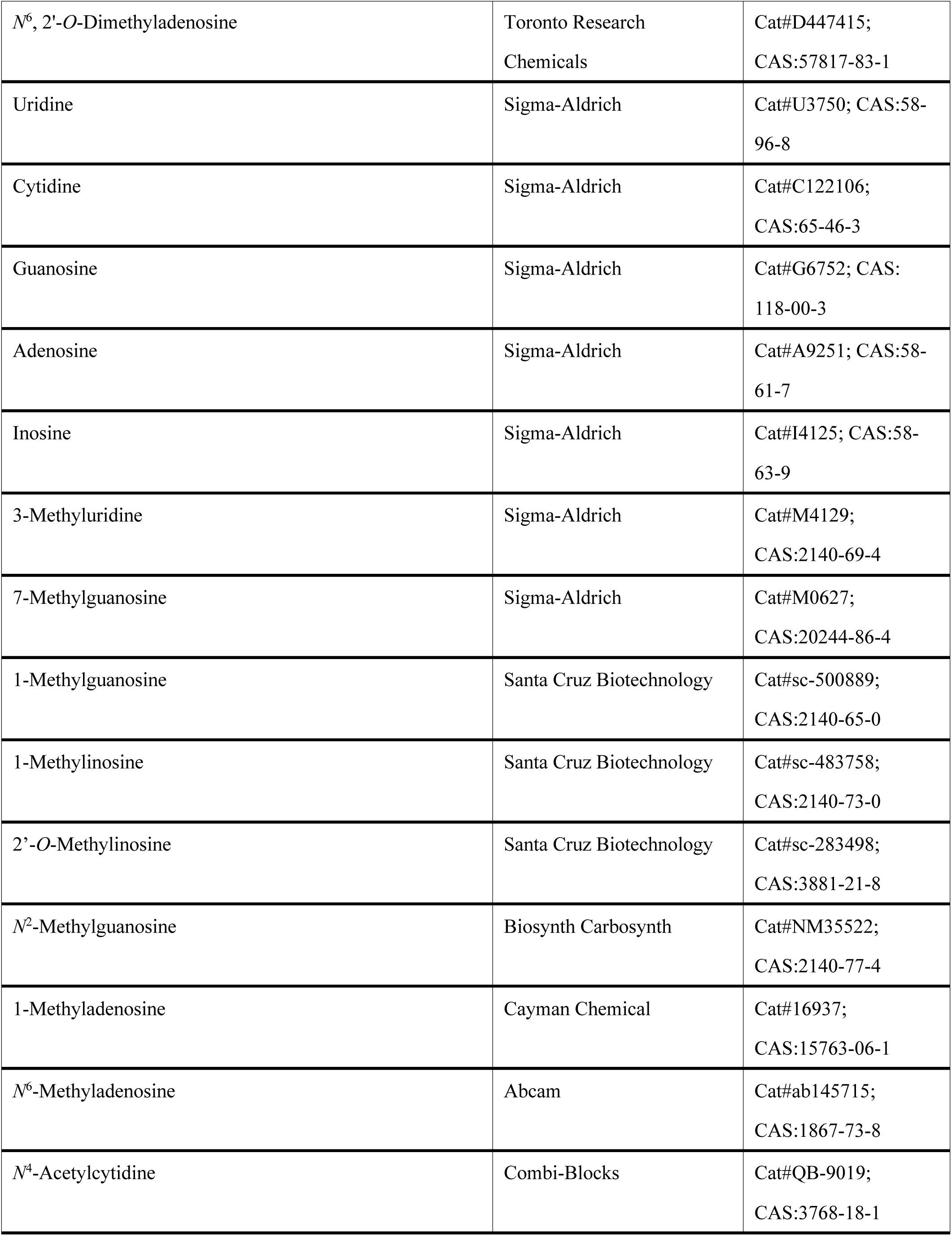

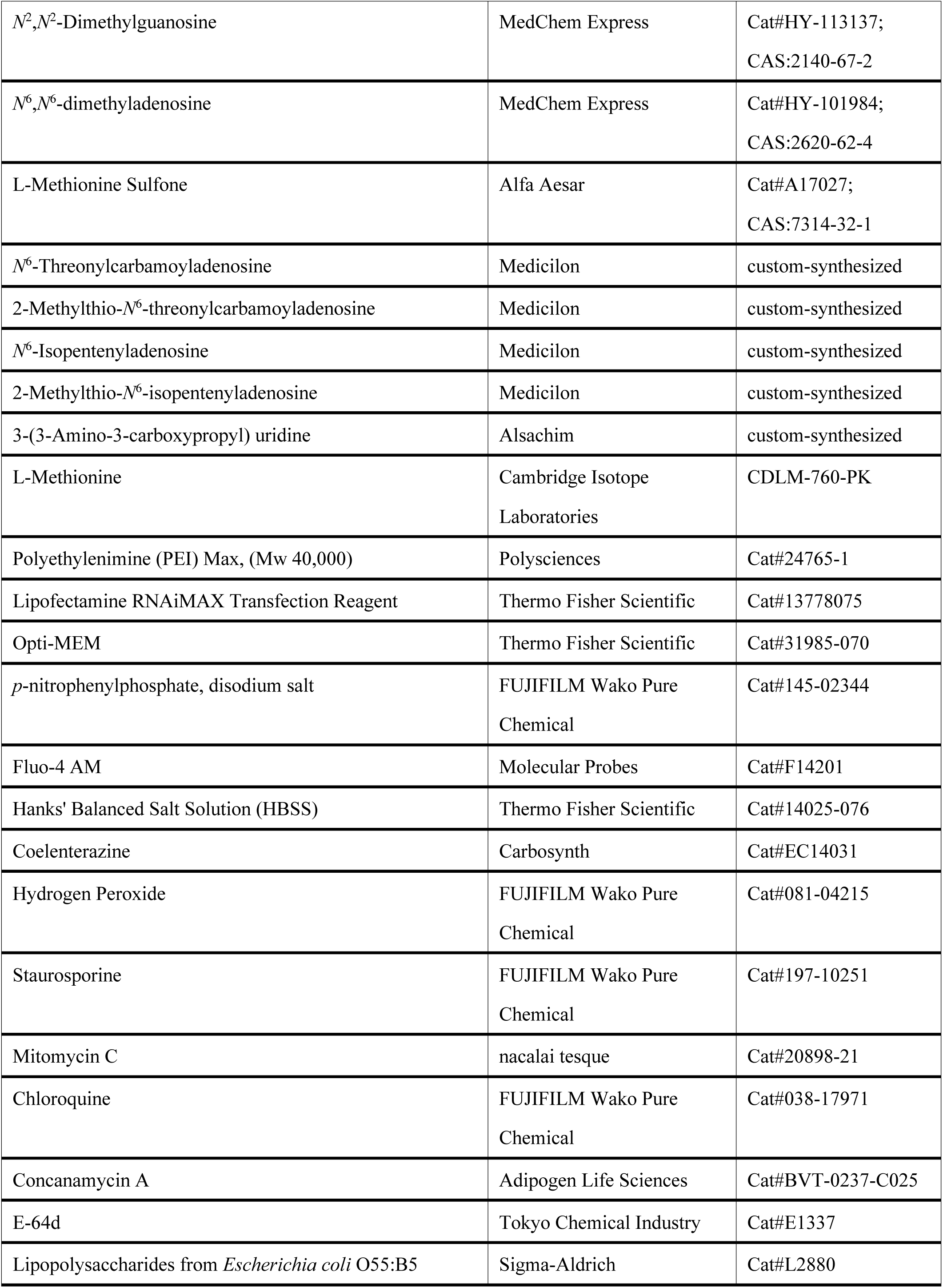

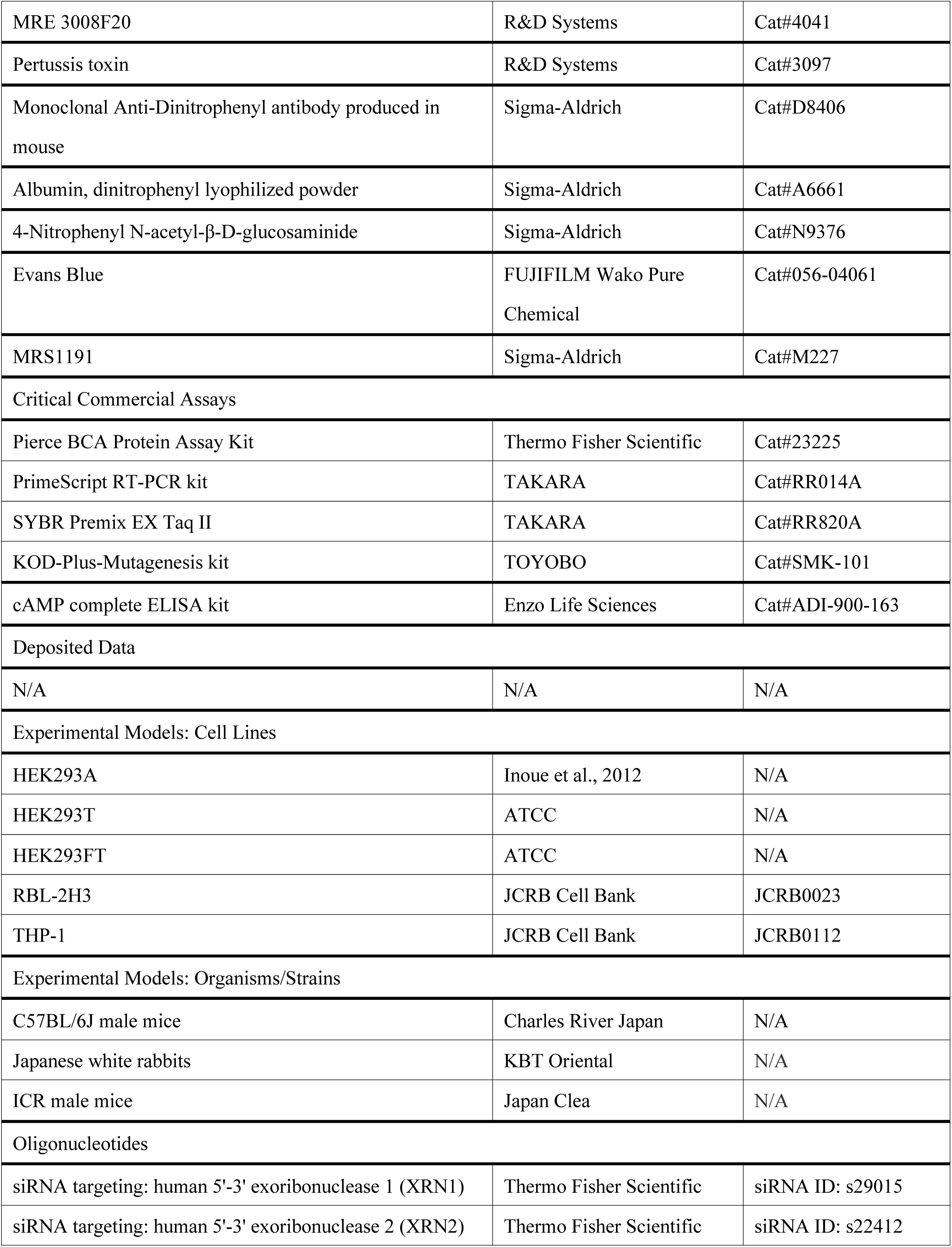

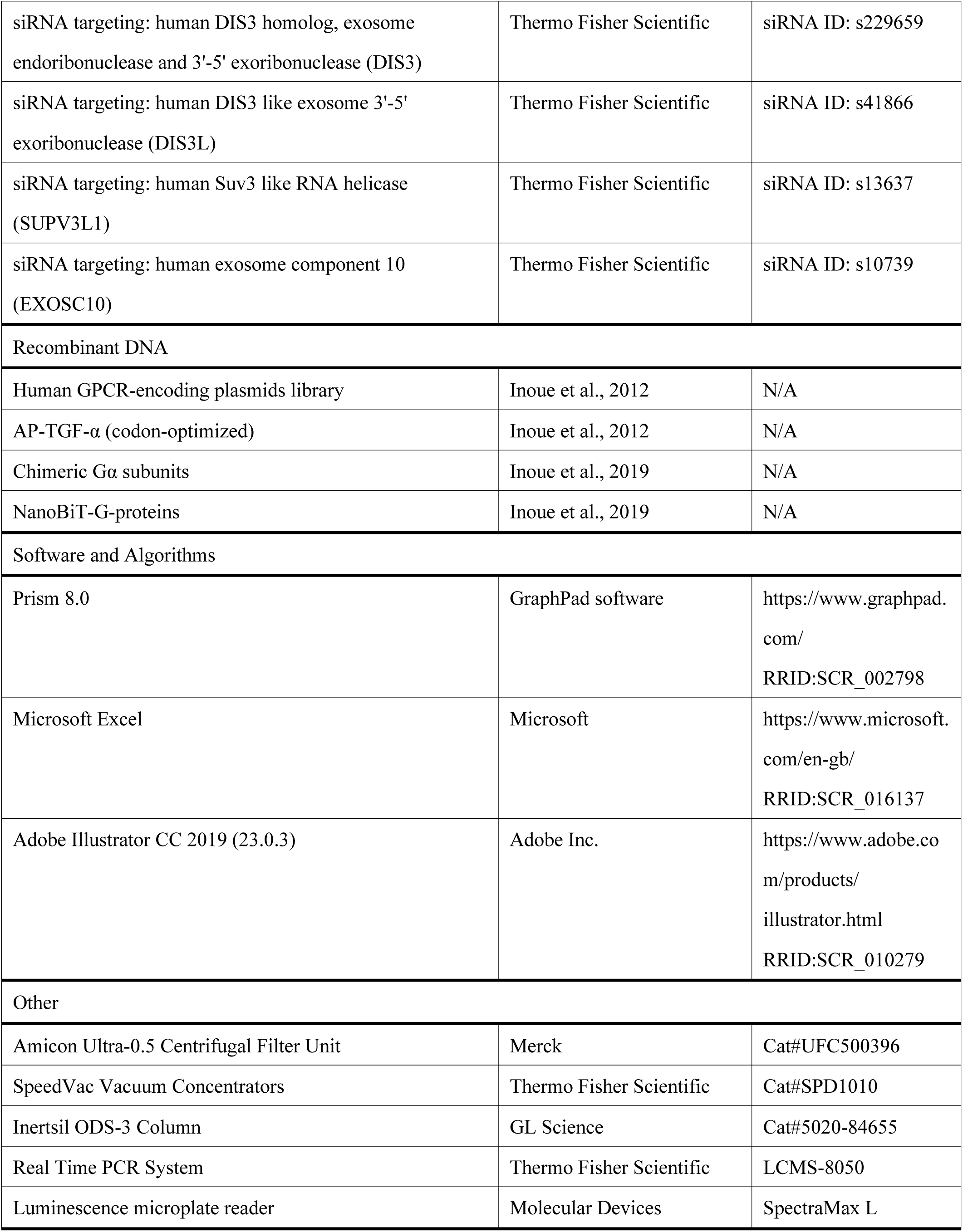

## METHODS DETAILS

### Biological sample collection and preparation for mass spectrometry analysis

Human plasma and serum samples (seven and six, respectively) collected from healthy donors older than 18 years of age were purchased from HemaCare Corporation. Human aqueous and vitreous humor samples (nine and two, respectively) were obtained from patients undergoing intraocular surgery for a macular hole at Kumamoto University Hospital. Twelve human urine samples were collected from healthy volunteers at Kumamoto University. These studies were approved by the Kumamoto University Ethics Committee (authorization no. ethics 1622 and ethics 1518), and written forms of informed consent were obtained from all subjects.

Experiments using animals were performed in accordance with the Declaration of Helsinki and the guidelines of the ARVO Statement for the Use of Animals in Ophthalmic and Vision Research, and were approved by the Kumamoto University Ethics Committee for Animal Experiments (authorization nos. A27-037 and D27-210). C57BL/6J male mice (6-8 weeks old) were purchased from Charles River Japan, and Japanese white rabbits (3 months old) were purchased from KBT Oriental. Mouse blood and urine, and rabbit aqueous humor, vitreous humor, and blood, were collected for the mass spectrometry analysis. Blood samples were allowed to clot at 4°C for 1 h and then centrifuged to obtain serum that were stored in aliquots at −80°C until analysis. Other biological samples were immediately frozen at −80°C until further analysis.

Metabolites in extracellular fluids of biological samples were extracted with ice-cold methanol/chloroform/ultrapure water containing 1 µM LMS as internal control. The suspension was then centrifuged at 16,000 *g* for 3 min at 4°C. Ultrapure water was added to the aqueous phase, and the suspension was centrifuged at 16,000 *g* for 3 min at 4°C. After centrifugation, the aqueous phase was ultrafiltered using an ultrafiltration tube (Amicon Ultra-0.5, 3 kDa; Merck Millipore). The filtrate was concentrated with a vacuum concentrator (SpeedVac; Thermo Fisher Scientific), and the concentrated filtrate was dissolved in 25 µl ultrapure water and subjected to LCMS analysis.

### Quantitative modified nucleoside analysis

Quantitative modified nucleoside analysis was performed using a triple quadrupole mass spectrometry system (LCMS-8050, Shimadzu) equipped with an electrospray ionization (ESI) source and an ultra-high performance liquid chromatography system^6^. Samples were injected into an Inertsil ODS-3 column (GL Science). The mobile phase consisted of 5 mM ammonium acetate (A), pH 5.3 in water and 60% acetonitrile in water (B). The LC gradient was set as follows: 1-10 min: 1-22.1% B, 10-15 min: 22.1-63.1% B, 15-17 min: 63.1-100% B, 17-22 min: 100% B, and 22-23 min, 100-0.6% B. The flow rate and injection volume were set at 0.4 ml/min and 2 μl, respectively. Detection was performed in multiple reaction monitoring (MRM) mode in LabSolutions System (Shimadzu). The MRM transitions for all nucleoside derivatives analyzed in this method are described in Supplementary Table 1. Interface temperature was 300°C, desolvation line temperature was 250°C, and heat block temperature was 400°C. Nitrogen gas was supplied by an N_2_ supplier Model T24FD (System Instruments) for nebulization and drying, and argon gas was used for collision-induced dissociation. Calibration curves using various concentrations of the nucleosides to be measured were obtained in every analytical run and then used to calculate the nucleoside concentrations of the samples.

### Cell culture and treatments

All cells were cultured at 37°C in an atmosphere of 5% CO_2_ unless stated otherwise. HEK293A, HEK293T, and HEK293FT cells were cultured in Dulbecco’s modified Eagle’s medium (DMEM, Gibco) supplemented with 10% heat-inactivated fetal bovine serum (FBS, Gibco). RBL-2H3 cells and THP-1 cells were cultured in Eagle’s minimal essential medium (EMEM, Wako) and Roswell Park Memorial Institute (RPMI) 1640 medium (Gibco), respectively, supplemented with heat-inactivated 10% FBS. Transient transfections were performed using either Lipofectamine RNAiMAX (Thermo Fisher) or polyethylenimine (PEI) transfection reagent (Polysciences). Transfection efficiency was analyzed by qPCR.

### TGF-α shedding assay

The transforming growth factor-α (TGFα) shedding assay, which measures the activation of GPCR receptor, was performed as described previously^20^. Briefly, pCAG plasmids encoding human A1R, A2AR, A2BR, or A3R construct were prepared. The A3R mutants I253T, V169E, L264T, and L264M; the A1R mutants T257I, E172V, and T270L; and the A2AR mutants T256I, E169V, and A2AR M270L were generated by introducing single-point mutations using the KOD-Plus-Mutagenesis kit. cDNAs for A3R of cow, pig, rat, mouse, chicken, frog, lizard, and fugu were synthesized and cloned into pCAG vector. HEK293A cells were seeded in a 6-well culture plates and then transfected with these plasmids, together with the plasmids encoding alkaline phosphatase (AP)-tagged TGFα (AP-TGFα) and chimeric Gα subunit proteins (Gα_q/o_ for A1R, Gα_q/s_ for A2AR and A2BR, and Gα_q/i3_ for A3R). After a 24 h culture, the transfected cells were harvested by trypsinization, neutralized with DMEM/FBS, and collected by centrifugation. Cells were suspended in Hank’s Balanced Salt Solution (HBSS, Gibco) containing 5 mM HEPES (pH 7.4) and were left for 10 min to remove extra AP-TGFα released during trypsinization. After centrifugation, cells were resuspended in HEPES-HBSS, and seeded in 96-well plates. The plates were incubated for 30 min to allow the cells to attach. Test compounds were diluted in 0.01% bovine serum albumin (BSA)-containing HEPES-HBSS and added to the cells. After a 1 h incubation, the 96-well plates were centrifuged, and the conditioned media was transferred to empty 96-well plates. AP reaction solution (a mixture of 10 mM p-nitrophenylphosphate (p-NPP), 120 mM Tris-HCl (pH 9.5), 40 mM NaCl, and 10 mM MgCl_2_) was added to plates containing cells and conditioned media. Absorbance at 405 nm was measured on a microplate reader (TECAN) before and after a 1 h incubation of the plates at room temperature. AP-TGFα release was calculated as described previously^20^. The AP-TGFα release percentages were fitted to a four-parameter sigmoidal concentration-response curve using the Prism 8 software (GraphPad), and the EC_50_ and E_max_ values were obtained. Receptor activation was scored using relative intrinsic activity (RAi)^21, 22^, which was defined as a relative E_max_/EC_50_ value.

### NanoBiT assay

The NanoBiT assays, which detects β-arrestin recruitment and G-protein dissociation, was performed as described previously^21, 54^. Briefly, HEK293A cells were seeded in 6-well plates and transfected with the indicated plasmids using the PEI transfection reagent. LgBiT-β-arrestin 2 (EE) and A3R-SmBiT expression vectors were used for β-arrestin recruitment assay, and GNAI1-LgBiT, GNB1, SmBiT-GNG2 (CS), and A3R expression vectors were used for G-protein dissociation assay. After a 24 h culture, the transfected cells were harvested and collected by 0.53 mM EDTA-containing D-PBS. After centrifugation, cells were resuspended in HEPES-HBSS containing 0.01% BSA and were seeded in 96-well plates. Coelenterazine was added to the plates at a final concentration of 10 µM, followed by incubation at room temperature for 2 h in the dark. After measurement of baseline luminescence using the Spectra Max L Microplate Reader (Molecular Devices), m^6^A or adenosine diluted in 0.01% BSA-HBSS was added. Kinetic luminescence was measured every 20 sec after compound addition. The average luminescence signal over 5-10 min was normalized against the initial value and was fitted to a four-parameter sigmoidal concentration-response curve. EC_50_ values were calculated using the Prism 8 software.

### Cytotoxic assays

To induce cytotoxicity, HEK293A cells were treated for 24 h with 100 μM, 500 μM, or 1 mM H_2_O_2_; 1 nM, 10 nM, or 100 nM staurosporine (Wako); or 0.0001%, 0.0005%, or 0.001% mitomycin C (MMC). For the hypoxia assay, THP-1 cells were maintained at either 21% O_2_ or 1% O_2_ for 48 h. Supernatants were collected, centrifuged, filtered (0.22 μm, Merck), and frozen at −80°C for subsequent analysis. For lysosomal inhibition, chloroquine (CLQ; Wako), concanamycin A (concA; Adipogen Life Sciences), and E-64d (Tokyo Chemical Industry) was applied to HEK 293A cells 12 h before the addition of 1 mM H_2_O_2_. Culture media was desalted with a C18 desalting column (GL science) and concentrated with a vacuum concentrator. The concentrated filtrate was dissolved in 25 μl ultrapure water and used for LCMS analysis.

### Generation of knockout cells by CRISPR-Cas9

Single-guide RNAs (sgRNA) were designed to target the human *ZCCHC4* or *METTL5* genomic locus. The sgRNA sequence driven by the U6 promoter was cloned into a lentiCRISPRv2 vector that also expresses Cas9 as previously described^55^. Lentiviral particles were produced in HEK293FT cells by co-transfection of the targeting vector with vectors expressing Gag, Pol, Rev, Tat, and VSV-G genes. The lentiviral plasmid DNA was then packed into lentivirus for infection in HEK293T cells. Infected cells were selected in puromycin for 2 weeks before single colonies were chosen and tested by DNA sequencing. For METTL5/ZCCHC4 double-knockout HEK cells, the METTL5-targeted sgRNA was cloned into a lentiGuide-Hygro-dTomato vector. The targeting vector or lentiCas9-Blast harboring Cas9 was transfected with the lentiviral expression system in HEK293FT cells for virus packaging. Supernatants were collected and used to infect *ZCCHC4*-knockout HEK293T cells. Infected cells were selected in hygromycin and blasticidin for 2 weeks before single colonies were chosen and tested by DNA sequencing.

### LPS-induced shock model of murine

C57BL/6J male mice were randomly divided into two groups of 12 mice each. Two groups were intraperitoneally injected with 20 mg/kg LPS (*Escherichia coli* 055: B5; Sigma-Aldrich) or vehicle (pyrogen-free saline). Blood and organ samples including liver, spleen and lung were collected 0, 6, and 12 h after LPS challenge. Serum was prepared from blood samples, and organ samples were immediately frozen in liquid nitrogen and stored at −80°C until lysate preparation. Lysates for quantification by LCMS were prepared as described above.

### Western blot analysis

For blots of pERK and ERK, A3R-transfected HEK293A cells were stimulated with 100 nM m^6^A for 5 min. Before stimulation with m^6^A, cells were pretreated with 1 μM MRE 3008F20 (R&D Systems) for 10 min or 150 ng/mL pertussis toxin (PTX; R&D Systems) overnight. Whole-cell lysates were prepared in RIPA Lysis Buffer (Thermo Fischer Scientific) containing a protease inhibitors (Thermo Fischer Scientific) and a phosphatase inhibitors (Nacalai Tesque). Cell lysates were sonicated and clarified by centrifugation. Protein concentrations were measured using the BCA Protein Assay kit (Pierce). Total protein was resuspended in NuPAGE LDS sample buffer. The same amount of protein was separated by SDS-PAGE (4-12% NuPAGE Bis-Tris gels; Invitrogen) in MOPS buffer (Life Technologies). Protein was transferred to 0.45 μm polyvinylidene difluoride (PVDF) membranes (Amersham). Membranes were blocked in 2% Block Ace (KAC), washed with Tris-buffered saline containing 0.1% Tween-20 (TBS-T), and incubated with the primary and corresponding secondary antibodies. Chemiluminescence signals were recorded with a luminescence image analyzer (ImageQuant LAS-4000, Fujifilm). Rabbit antibodies against human p-ERK1/2 and ERK were purchased from Cell Signaling.

### Intracellular calcium imaging

HEK293 cells or RBL-2H3 cells were loaded with 2.5 μM Fluo-4 AM (Molecular Probes) for 30 min, and intracellular calcium transients were recorded every 5 sec using a confocal microscope (Olympus FV 3000). After the basal calcium level was monitored for 2 min, m^6^A was applied to monitor calcium dynamics. MRE 3008F20 (10 µM for HEK293 cells and 1 µM for RBL-2H3 cells) was applied 10 min before the addition of m^6^A (10 µM for HEK 293 cells and 5 µM for RBL-2H3 cells). Signals from ten representative cells were digitized using the Fluoview software (Olympus), and the data were summarized in a graph using the Prism 8 software.

### Degranulation assay

The degranulation response of the RBL-2H3 basophils was quantified by measuring the levels of β-hexosaminidase released into the supernatants as previously described with slight modification^56^. Briefly, RBL-2H3 cells were seeded in 96-well plates, incubated for 24 h, and then sensitized for 12 h with 1 μg/ml anti-DNP-IgE (Sigma-Aldrich). After washing four times with HBSS, the cells were exposed to different concentrations of m^6^A (0–5000 nM) for 1 h, and then stimulated with 100 ng/mL DNP-BSA for 1 h. The supernatant was transferred to an empty 96-well plate, and the cells were lysed with 1% Triton X-100/PBS. Supernatants and cell lysates were incubated with 1 mM p-nitrophenyl-N-acetyl-β-D-glucosaminide in 0.05 M citrate buffer (pH 4.5) at 37°C for 1 h. The reaction was terminated by addition of 0.05 M sodium carbonate buffer (pH 10.0). p-nitrophenol, the product of the reaction, was detected by measuring optical absorbance at 405 nm.

### cAMP quantification

RBL-2H3 cells seeded in 12-well plates were stimulated with forskolin (0.5 μM) and the indicated concentrations of m^6^A (0–1000 nM). Intracellular cAMP level was quantified using an ELISA kit (Enzo Life Sciences).

### Passive cutaneous anaphylaxis (PCA) mouse model

PCA reactions in mice, used as an animal model for type 1 allergy, were performed as previously described with slight modifications^36^. In brief, ICR male mice (Japan Clea) were administrated intradermally with 100 ng anti-DNP IgE into the ear. After 24 h, the IgE-sensitized ears were injected intradermally with different concentrations of m^6^A (0-1.0 mg/mL), and 5 min later the mice were challenged with an intravenous injection of 50 μg DNP-BSA containing 0.5 % Evans Blue dye (Wako). The mice were euthanized 5 min after the challenge, and the treated ear was excised to measure the amount of dye extravasated. Dye was extracted from the ear in 700 μl formamide at 56 °C overnight, and absorbance was measured at 620 nm. In experiments using A3R antagonist, 0.01 mg/mL MRS 1191 (Sigma-Aldrich) and 0.1 mg/mL m^6^A were administrated simultaneously.

### Modeling of the m^6^A bound A3 receptor

The structure of the human adenosine A3 receptor was prepared by the web-server SWISS-MODEL^57, 58^, using the Gi-complexed human adenosine A1 receptor as a template. m^6^A was modeled manually, based on the coordination of adenosine in the A1 receptor (PDB code: 6D9H)

### Statistics and reproducibility

Representative results from at least two independent experiments are shown for every figure, unless stated otherwise in the figure legends.

## DATA AVAILABILITY

The authors declare that the data supporting the findings of this study are available within the paper and its supplementary information files. Data not included are available from the corresponding authors upon reasonable request.

## Notes

### Competing Interest Statement

The authors have declared no competing interest.

## REFERENCES

Arsenis, C., Gordon, J. S., and Touster, O. (1970). Degradation of nucleic acids by lysosomal extracts of rat liver and Ehrlich ascites tumor cells. J. Biol. Chem. 245, 205–211.

Biasini, M., Bienert, S., Waterhouse, A., Arnold, K., Studer, G., Schmidt, T., Kiefer, F., Gallo Cassarino, T., Bertoni, M., Bordoli, L., et al. (2014). SWISS-MODEL: modelling protein tertiary and quaternary structure using evolutionary information. Nucleic. Acids. Res. 42, W252–258.

Boccaletto, P., Machnicka, M. A., Purta, E., Piatkowski, P., Baginski, B., Wirecki, T. K., de Crécy-Lagard, V., Ross, R., Limbach, P. A., Kotter, A., et al. (2018) MODOMICS: a database of RNA modification pathways. 2017 update. Nucleic. Acids. Res. 46, D303-D307.

Borea, P. A., Gessi, S., Merighi, S., Vincenzi, F., and Varani K. (2018). Pharmacology of Adenosine Receptors: The State of the Art. Physiol Rev. 98, 1591–1625.

Borea, P. A., Varani, K., Vincenzi, F., Baraldi, P. G., Tabrizi, M.A., Merighi, S., and Gessi, S. (2015) The A3 adenosine receptor: history and perspectives. Pharmacol. Rev. 67, 74–102.

Borek, E., Baliga, B. S., Gehrke, C. W., Kuo, C. W., Belman, S., Troll, W., and Waalkes, T. P. (1977). High turnover rate of transfer RNA in tumor tissue. Cancer. Res. 37, 3362–3366.

Carpenter, B., Nehmé, R., Warne, T., Leslie, A. G., and Tate, C. G. (2016). Structure of the adenosine A2A receptor bound to an engineered G protein. Nature 536, 104–107.

Chen, M., Urs, M. J., Sánchez-González, I., Olayioye, M. A., Herde, M., and Witte, C. P. (2018). m^6^A RNA Degradation Products Are Catabolized by an Evolutionarily Conserved N^6^-Methyl-AMP Deaminase in Plant and Mammalian Cells. Plant Cell. 30, 1511–1522.

Cheng, R. K. Y., Segala, E., Robertson, N., Deflorian, F., Doré, A. S., Errey, J. C., Fiez-Vandal, C., Marshall, F. H., and Cooke, R. M. (2017). Structures of Human A1 and A2A Adenosine Receptors with Xanthines Reveal Determinants of Selectivity. Structure 25, 1275–1285.

De Jesus, D. F., Zhang, Z., Kahraman, S., Brown, N. K., Chen, M., Hu, J., Gupta, M. K., He, C., and Kulkarni, R. N. (2019). m^6^A mRNA Methylation Regulates Human β-Cell Biology in Physiological States and in Type 2 Diabetes. Nat Metab. 1,765–774.

Draper-Joyce, C. J., Khoshouei, M., Thal, D. M., Liang, Y. L., Nguyen, A. T. N., Furness, S. G. B., Venugopal, H., Baltos, J. A., Plitzko, J. M., Danev, R., et. al. (2018). Structure of the adenosine-bound human adenosine A1 receptor-Gi complex. Nature 558, 559–563.

Ehlert, F. J., Griffin, M. T., Sawyer, G. W., and Bailon, R. (1999). A simple method for estimation of agonist activity at receptor subtypes: comparison of native and cloned M3 muscarinic receptors in guinea pig ileum and transfected cells. J. Pharmacol. Exp. Ther. 289, 981–992.

Eltzschig, H. K. (2009). Adenosine: An Old Drug Newly Discovered. Anesthesiology. 111, 904–915.

Fakruddin, M., Wei, F. Y., Suzuki, T., Asano, K., Kaieda, T., Omori, A., Izumi, R., Fujimura, A., Kaitsuka, T., Miyata, K., et al. (2018). Defective Mitochondrial tRNA Taurine Modification Activates Global Proteostress and Leads to Mitochondrial Disease. Cell Rep. 22, 482–496.

Fiala, E. S., Conaway, C. C., and Mathis, J. E. (1989). Oxidative DNA and RNA damage in the livers of Sprague Dawley rats treated with the hepatocarcinogen 2-nitropropane. Cancer Res. 49, 5518–5522.

Frye, M., Jaffrey, S. R., Pan, T., Rechavi, G., and Suzuki, T. (2016). RNA modifications: what have we learned and where are we headed? Nat. Rev. Genet. 17, 365–372.

Fujiwara, Y., Furuta, A., Kikuchi, H., Aizawa, S., Hatanaka, Y., Konya, C., Uchida, K., Yoshimura, A., Tamai, Y., Wada, K., et al. (2013). Discovery of a novel type of autophagy targeting RNA. Autophagy 9, 403–409.

Gilbert, W. V., Bell, T. A., and Schaening, C. (2016). Messenger RNA modifications: form, distribution, and function. Science 352, 1408–1412.

Gmitterová, K., Gawinecka, J., Heinemann, U., Valkovič, P., and Zerr, I. (2018). DNA versus RNA oxidation in Parkinson’s disease: Which is more important? Neurosci Lett. 662, 22–28.

Hisano, Y., Kono, M., Cartier, A., Engelbrecht, E., Kano, K., Kawakami, K., Xiong, Y., Piao, W., Galvani, S., Yanagida, K., et al. (2019). Lysolipid receptor cross-talk regulates lymphatic endothelial junctions in lymph nodes. J Exp Med. 216, 1582–1598.

Houseley, J., and Tollervey, D. (2009). The many pathways of RNA degradation. Cell 136, 763–776.

Inoue, A., Raimondi, F., Kadji, F. M.N., Singh, G., Kishi, T., Uwamizu, A., Ono, Y., Shinjo, Y., Ishida, S., Arang, N., et al. (2019). Illuminating G-Protein-Coupling Selectivity of GPCRs. Cell 177, 1933–1947.

Inoue, A., Ishiguro, J., Kitamura, H., Arima, N., Okutani, M., Shuto, A., Higashiyama, S., Ohwada, T., Arai, H., Makide, K., et al. (2012). TGFα shedding assay: an accurate and versatile method for detecting GPCR activation. Nat. Methods. 9, 1021–1029.

Jackson, E. K., Kotermanski, S. E., Menshikova, E. V., Dubey, R. K., Jackson, T. C., and Kochanek, P. M. (2017). Adenosine production by brain cells. J. Neurochem. 141, 676–693.

Jacobson, K. A., and Gao, Z. G. (2006). Adenosine receptors as therapeutic targets. Nat. Rev. Drug. Discov. 5, 247–264.

Jonkhout, N., Tran, J., Smith, M. A., Schonrock, N., Mattick, J. S., and Novoa, E. M. (2017). The RNA modification landscape in human disease. RNA. 12, 1754–1769.

Lan, Q., Liu, P. Y., Haase, J., Bell, J. L., Hüttelmaier, S., and Liu, T. (2019). The Critical Role of RNA m^6^A Methylation in Cancer. Cancer. Res. 79, 1285–1292.

Liu, N., Zhou, K. I., Parisien, M., Dai, Q., Diatchenko, L., and Pan, T. (2017). N6-methyladenosine alters RNA structure to regulate binding of a low-complexity protein. Nucleic Acids Res. 45, 6051–6063.

Lothrop, C. D. Jr., and Uziel, M. (1982). Salvage of the modified nucleoside ribothymidine in cultured hamster embryo cells. Biochim Biophys Acta. 698, 134–139 (1982)

Ma, H., Wang, X., Cai, J., Dai, Q., Natchiar, S. K., Lv, R., Chen, K., Lu, Z., Chen, H., Shi, Y. G., et al. (2019). N6-Methyladenosine methyltransferase ZCCHC4 mediates ribosomal RNA methylation. Nat. Chem. Biol. 15, 88–94.

Mandel, L. R., Srinivasan, P. R., and Borek, E. (1966). Origin of urinary methylated purines. Nature 209, 586–588.

Nunomura, A., Perry, G., Pappolla, M. A., Wade, R., Hirai, K., Chiba, S., and Smith, M. A. (1999). RNA oxidation is a prominent feature of vulnerable neurons in Alzheimer’s disease. J Neurosci. 19, 1959–1964.

Ortega, E., Hazan, B., Zor, U., and Pecht, I. (1989). Mast cell stimulation by monoclonal antibodies specific for the Fc epsilon receptor yields distinct responses of arachidonic acid and leukotriene C4 secretion. Eur. J. Immunol. 19, 2251–2256.

Ovary, Z. (1958). Passive cutaneous anaphylaxis in the mouse. J. Immunol. 81, 355–357.

Pane, F., Oriani, G., Kuo, K. C., Gehrke, C. W., Salvatore, F., and Sacchetti, L. (1992). Reference intervals for eight modified nucleosides in serum in a healthy population from Italy and the United States. Clin. Chem. 38, 671–677.

Reeves, J. J., Jones, C. A., Sheehan, M. J., Vardey, C. J., and Whelan, C. J. (1997). Adenosine A3 receptors promote degranulation of rat mast cells both in vitro and in vivo. Inflamm. Res. 46, 180–184.

Roundtree, I. A., Evans, M. E., Pan, T., and He, C. (2017a). Dynamic RNA Modifications in Gene Expression Regulation. Cell 169, 1187–1200.

Roundtree, I. A., Luo, G. Z., Zhang, Z., Wang, X., Zhou, T., Cui, Y., Sha, J., Huang, X., Guerrero, L., Xie, P., et al. (2017b). YTHDC1 mediates nuclear export of N^6^-methyladenosine methylated mRNAs. Elife. 6, e31311.

Saito, H., Nishimura, M., Shinano, H., Makita, H., Tsujino, I., Shibuya, E., Sato, F., Miyamoto, K., Kawakami, Y., et al. (1999). Plasma concentration of adenosine during normoxia and moderate hypoxia in humans. Am. J. Respir. Crit. Care. Med. 159, 1014–1018.

Schwede, T., Kopp, J., Guex, N., and Peitsch, M. C. (2003). SWISS-MODEL: An automated protein homology-modeling server. Nucleic. Acids. Res. 31, 3381–3385.

Seidel, A., Brunner, S., Seidel, P., Fritz, G. I., and Herbarth, O. (2006). Modified nucleosides: an accurate tumour marker for clinical diagnosis of cancer, early detection and therapy control. Br J Cancer 94, 1726–1733.

Shalem, O., Sanjana, N. E., Hartenian, E., Shi, X., Scott, D. A., Mikkelson, T., Heckl, D., Ebert, B. L., Root, D. E., Doench, J. G., et al. (2014). Genome-scale CRISPR-Cas9 knockout screening in human cells. Science 343, 84–87.

Shi, H., Zhang, X., Weng, Y. L., Lu, Z., Liu, Y., Lu, Z., Li, J., Hao, P., Zhang, Y., Zhang, F. et al. (2018). m^6^A facilitates hippocampus-dependent learning and memory through YTHDF1. Nature 563, 249–253.

Shibata, T., Ohto, U., Nomura, S., Kibata, K., Motoi, Y., Zhang, Y., Murakami, Y., Fukui, R., Ishimoto, T., Sano, S. et al. (2016). Guanosine and its modified derivatives are endogenous ligands for TLR7. Int Immunol. 28, 211–222.

Shneyvays, V., Zinman, T., and Shainberg, A. (2004). Analysis of calcium responses mediated by the A3 adenosine receptor in cultured newborn rat cardiac myocytes. Cell Calcium 36, 387–396.

Uziel, M., and Selkirk, J. J. (1980). Pyrimidine nucleotide pool changes during cell cycle and quiescence. Pyrimidine excretion and metabolic isolation of the pyrimidine nucleoside polyphosphates. Biol. Chem, 255, 11227–11232.

Uziel, M., and Selkirk, J. J. (1979). Pyrimidine nucleoside, pseudouridine, and modified nucleoside excretion by growing and resting fibroblasts. Cell. Physiol. 99, 217–222.

van Tran, N., Ernst, F. G. M., Hawley, B. R., Zorbas, C., Ulryck, N., Hackert, P., Bohnsack, K. E., Bohnsack, M. T., Jaffrey, S. R., Graille, M., et al. (2019). The human 18S rRNA m6A methyltransferase METTL5 is stabilized by TRMT112. Nucleic. Acids. Res. 47, 7719–7733.

Wang, X., Zhao, B. S., Roundtree, I. A., Lu, Z., Han, D., Ma, H., Weng, X., Chen, K., Shi, H., and He, C. (2015). N(6)-methyladenosine Modulates Messenger RNA Translation Efficiency. Cell 161, 1388–1399.

Wang, X. Lu, Z., Gomez, A., Hon, G. C., Yue, Y., Han, D., Fu, Y., Parisien, M., Dai, Q., Jia, G., et al. (2014). N6-methyladenosine-dependent regulation of messenger RNA stability. Nature 505, 117–120.

Wei, F. Y., Zhou, B., Suzuki, T., Miyata, K., Ujihara, Y., Horiguchi, H., Takahashi, N., Xie, P., Michiue, H., Fujimura, A., et al. (2015). Cdk5rap1-mediated 2-methylthio modification of mitochondrial tRNAs governs protein translation and contributes to myopathy in mice and humans. Cell Metab. 21, 428–442.

Wei, F. Y., Suzuki, T., Watanabe, S., Kimura, S., Kaitsuka, T., Fujimura, A., Matsui, H., Atta, M., Michiue, H., Fontecave, M., et al. (2011). Deficit of tRNALys modification by Cdkal1 causes the development of type 2 diabetes in mice. J. Clin. Invest. 121, 3598– 3608.

Willmann, L., Erbes, T., Krieger, S., Trafkowski, J., Rodamer, M., and Kammerer, B. (2015). Metabolome analysis via comprehensive two-dimensional liquid chromatography: identification of modified nucleosides from RNA metabolism. Anal Bioanal Chem. 407, 3555–3566.

Winkler, R., Gillis, E., Lasman, L., Safra, M., Geula, S., Soyris, C., Nachshon, A., Tai-Schmiedel, J., Friedman, N., Le-Trilling, V. T. K., et al. (2019). m^6^A modification controls the innate immune response to infection by targeting type I interferons. Nat. Immunol. 20, 173–182.

Xiao, W., Adhikari, S., Dahal, U., Chen, Y. S., Hao, Y. J., Sun, B. F., Sun, H. Y., Li, A., Ping, X. L., Lai, W. Y., et al. (2016). Nuclear m(6)A Reader YTHDC1 Regulates mRNA Splicing. Mol. Cell 61, 507–519.

Zhao, B., Roundtree, I. A., and He, C. (2017). Post-transcriptional gene regulation by mRNA modifications. Nat. Rev. Mol. Cell Biol. 18, 31–42.

