## Supplemental Data for "*N*^6^-methyladenosine (m^6^A) is an endogenous A3 adenosine receptor ligand"

### Supplementary Figures

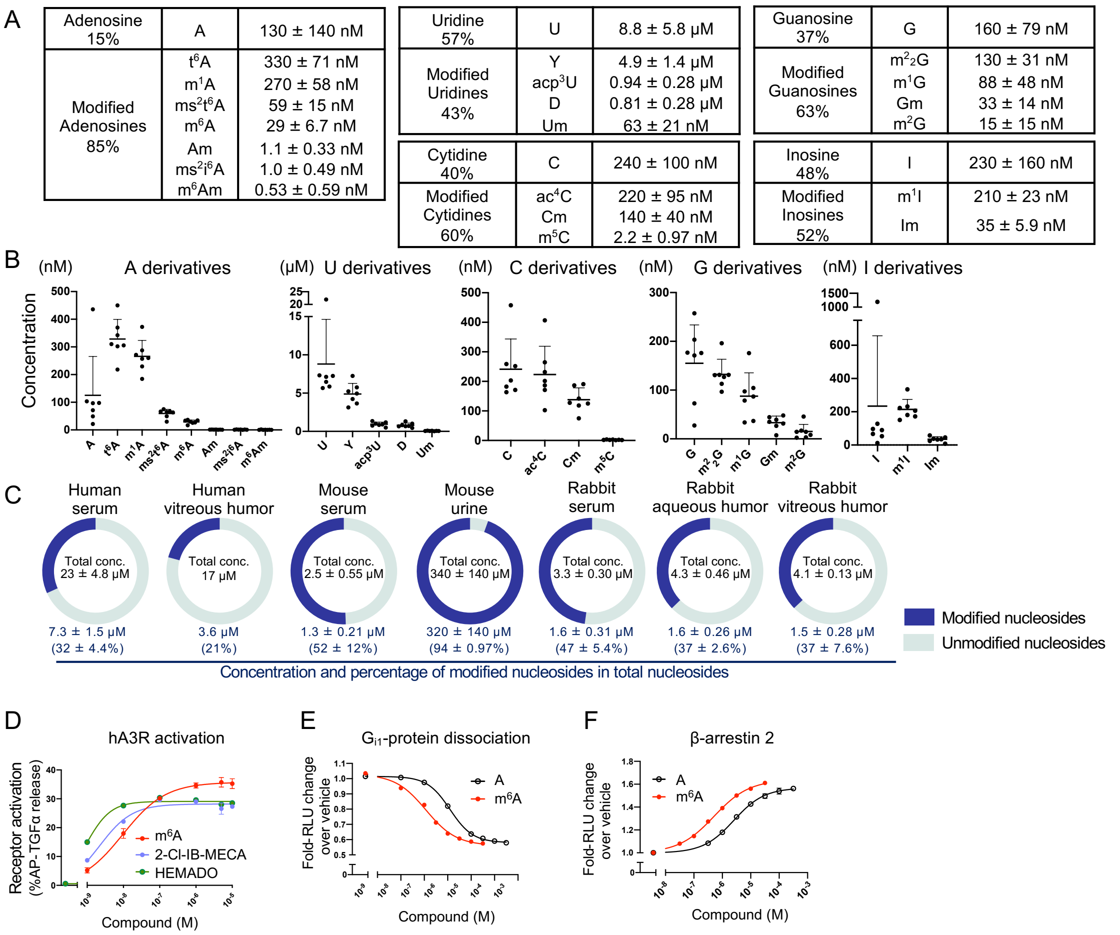


**Figure S1. Nucleoside distribution in biological extracellular fluid and m$^6$A activation of A3R, related to Figure 1.**

(A, B) Nucleoside derivatives in human plasma (n = 7) were grouped into subclasses. (A) Average concentrations of the nucleoside derivatives. Data are shown as means ± SD. (B) Individual values of nucleoside derivatives. Data are shown as means ± SD.

(C) Distribution of nucleoside derivatives in human serum (n = 6), human vitreous humor (n = 2), mouse serum (n = 6), mouse urine (n = 6), rabbit serum (n = 3), rabbit aqueous humor (n = 3), and rabbit vitreous humor (n = 3). Total concentration of nucleoside in each extracellular fluid is shown in the middle of the pie chart, and the concentration and percentage of modified nucleosides in total nucleosides are shown below the pie charts. Data are shown as mean ± SD.

(D) TGF-α shedding response curves of $m^6A$ and commercially available A3R synthetic agonists. Symbols and error bars represent means and SEM, respectively, of two independent experiments with each performed in triplicate.

(E) $m^6A$- or adenosine-mediated $G_{i1}$ dissociation was evaluated by NanoBiT-G-protein dissociation assay. $m^6A$ induced $G_{i1}$ dissociation of A3R more strongly than adenosine. Symbols are means of representative experiments with each performed in duplicate. Parameters (means ± SEM) obtained from three independent experiment are shown at the right.

(F) $m^6A$- or adenosine-mediated β-arrestin 2 recruitment was evaluated by NanoBiT assay. $m^6A$ induced β-arrestin recruitment to A3R more strongly than adenosine. Symbols are means of the representative experiments with each performed in duplicate. Parameters (means ± SEM) obtained from three independent experiment are shown to the right.

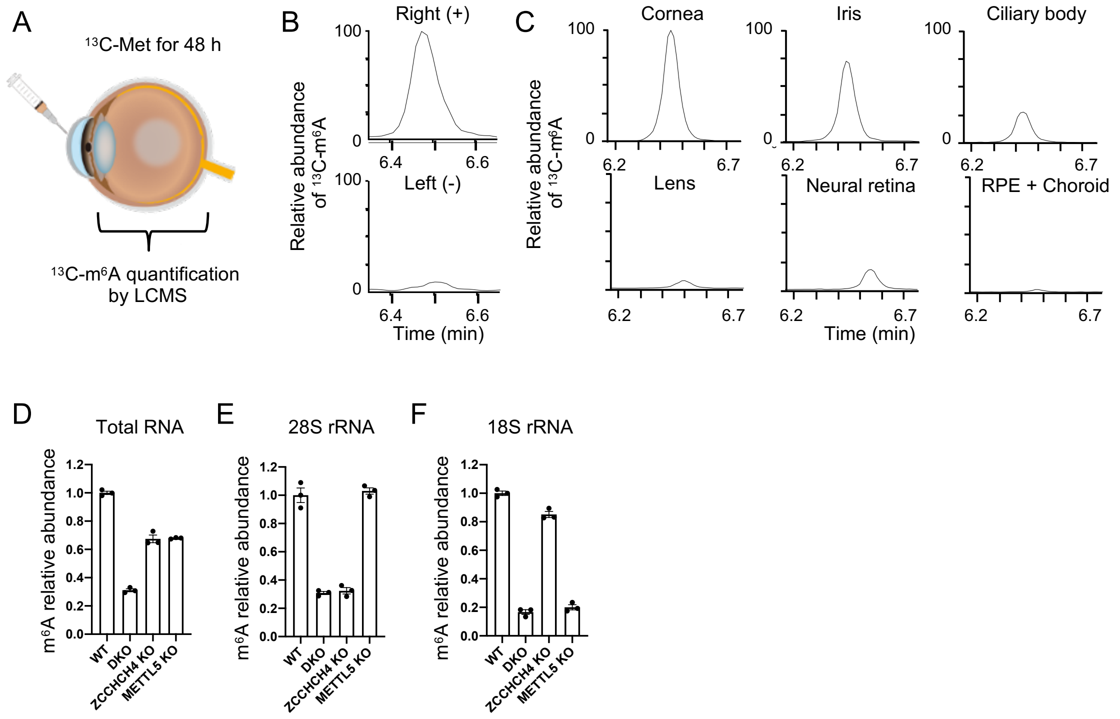


**Figure S2. Extracellular m$^6$A is derived from multiple RNA species, related to Figure 2.**

(A) Schematic of the m$^6$A synthesis detection *in vivo*.

(B and C) Aqueous humor (B) and ocular tissues (C) were collected 48 h after rabbits received an intracameral injection of stable isotope methionine only into the right eye. $^{13}$C-m$^6$A was detected in the aqueous humor of the right eye. At RNA levels, $^{13}$C-m$^6$A was mainly detected in the anterior part of the right eye, with the highest level in cornea (n = 3).

(D) $m^6A$ abundance in total RNA in the cells lacking $m^6A$ modifications in 28S and 18S rRNA. $m^6A$ level in total RNA decreased to 67% of wild-type levels in *ZCCHC4* KO cells, 68% in *METTL5* KO cells, and 31% in DKO cells. Bar graph is representative of two independent experiments (n = 3 each). Data are means ± SEM.

(E) $m^6A$ relative abundance in 28S rRNA. The $m^6A$ levels decreased to 34% and 32% of wild-type levels in *ZCCHC4* KO cells and DKO cells, respectively, but did not decrease in *METTL5* KO cells. Bar graph is representative of two independent experiments (n = 3 each). Data are means ± SEM.

(F) $m^6A$ relative abundance in 18S rRNA. The $m^6A$ level decreased to 21% of wild-type levels in *METTL5* KO cells and 14% in DKO cells. The $m^6A$ level decreased only mildly (to 86% of wild-type levels) in *ZCCHC4* KO cells. Bar graph is representative of two independent experiments (n = 3 each). Data are means ± SEM.

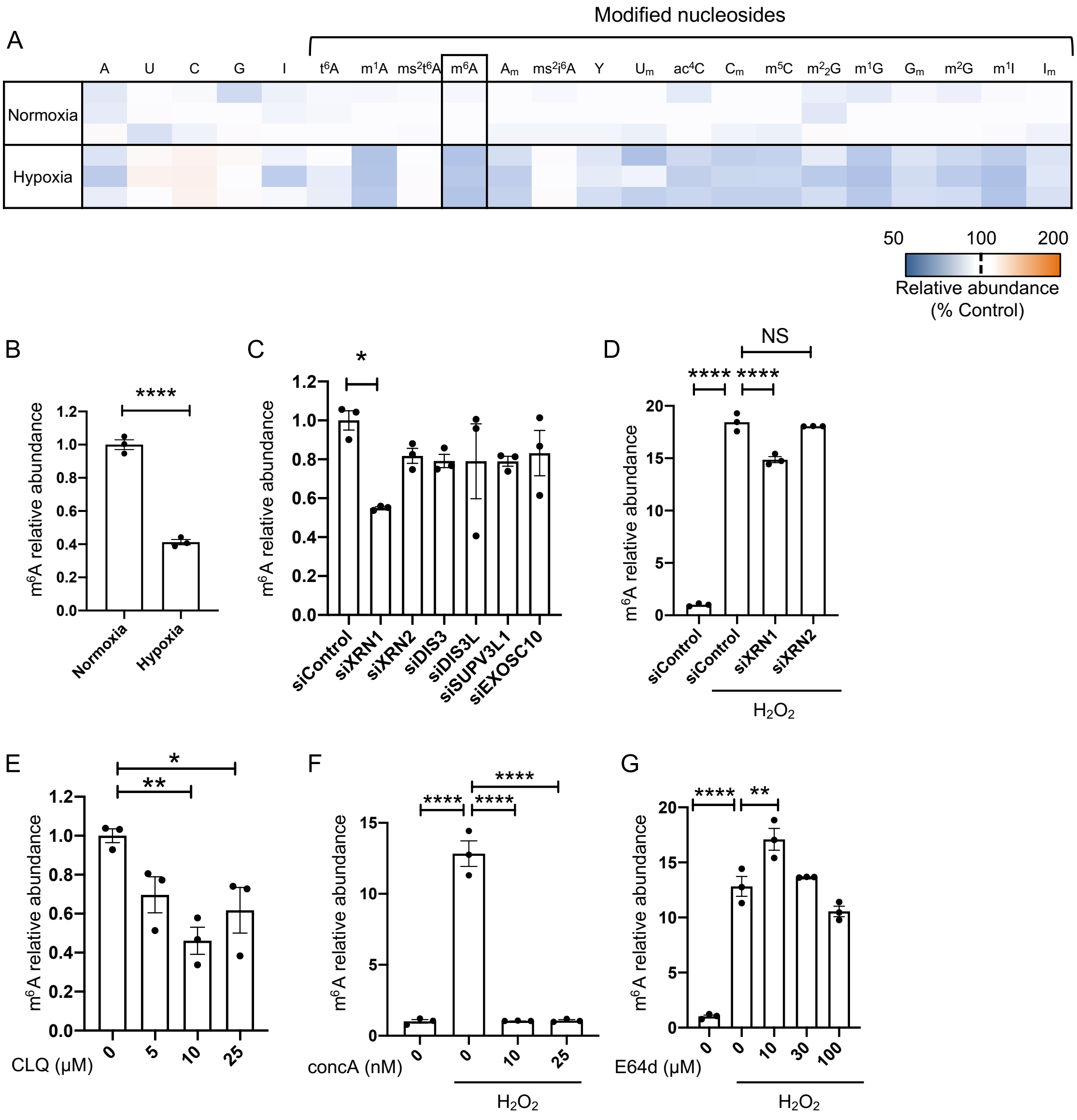


**Figure S3. The regulation of m$^6$A release, related to Figure 3.**

(A) Heatmap of extracellular nucleoside derivatives in supernatants of THP1 cells exposed to hypoxia. Two (n = 3) independent experiments were performed and the representative values are shown.

(B) Extracellular $m^6A$ levels in THP1 cells were reduced upon 48 h treatment with 1% $O_2$ hypoxia. ****P < 0.0001 in normoxia vs. hypoxia (unpaired two-tailed Student's *t*-test). Bar graph is representative of two independent experiments, three biological replicates each. Data are means ± SEM.

(C) Involvement of ribonucleases in extracellular $m^6A$ at steady state. HEK293A cells were treated with $^{13}C$-labeled methionine for 12 h after transfection with siRNAs targeting ribonucleases. Supernatants were collected 48 h after medium was replaced with normal medium, and $^{13}C$-$m^6A$ levels were evaluated. Among key ribonucleases, knockdown of 5'-3' exoribonuclease 1 (XRN1) decreased $m^6A$ levels by 45%. *P < 0.05 vs. siControl group (one-way ANOVA followed by a Dunnett's multiple comparison test). Bar graph is representative of two independent experiments (n = 3 each). Data are means ± SEM.

(D) Ribonuclease knockdown of XRN1 moderately suppressed $H_2O_2$-mediated $m^6A$ release. HEK293A cells were transfected with siRNAs, and the supernatants were collected 24 h after the addition of 1 mM $H_2O_2$. ****P < 0.0001, *P < 0.05 vs. siControl treated with 1 mM $H_2O_2$ (one-way ANOVA followed by a Dunnett's multiple comparison test). Bar graph is representative of three independent experiments (n = 2 each). Bar represents means.

(E) Involvement of lysosome in extracellular $m^6A$ release at steady state. HEK293A cells were treated with the indicated concentrations of chloroquine (CLQ) for 12 h, and supernatants were collected after 48 h. **P < 0.01 and *P < 0.05 vs. control group (one-way ANOVA followed by a Dunnett's multiple comparison test). Bar graph is representative of two independent experiments (n = 3 each). Data are means ± SEM.

(F and G) Involvement of lysosome in extracellular $m^6A$ release following $H_2O_2$-induced cytotoxicity. HEK293A cells were treated with the indicated concentrations of (F) concanamycin A (concA) or (G) E64d for 12 h, and the supernatants were collected 24 h after 1 mM $H_2O_2$ was added to the cells. **P < 0.01 vs. 1 mM $H_2O_2$ (one-way ANOVA followed by a Dunnett's multiple comparison test). Bar graph is representative of three independent experiments (n = 2 each). Bar represents means.

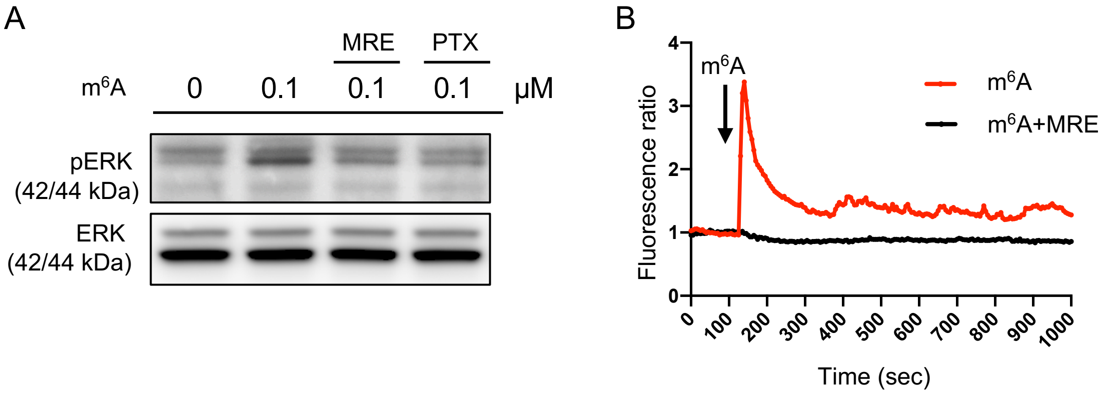


**Figure S4. Extracellular m$^6$A activates intracellular signaling transduction via A3R, related to Figure 4.**

(A) Representative western blot (n = 3) for m$^6$A-mediated phosphorylation of ERK in HEK cells overexpressing A3R.

(B) Intracellular calcium transient in HEK cells overexpressing A3R in response to 10 µM m$^6$A. m$^6$A-induced calcium transients were effectively blocked by pretreatment with the A3R antagonist (10 µM MRE 3008F20; MRE). Transients of ten representative cells are shown.

A

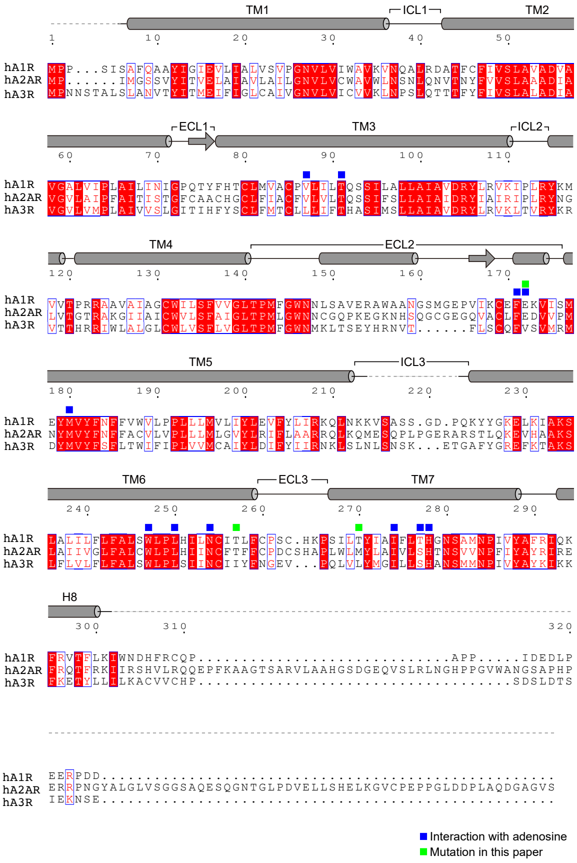


C

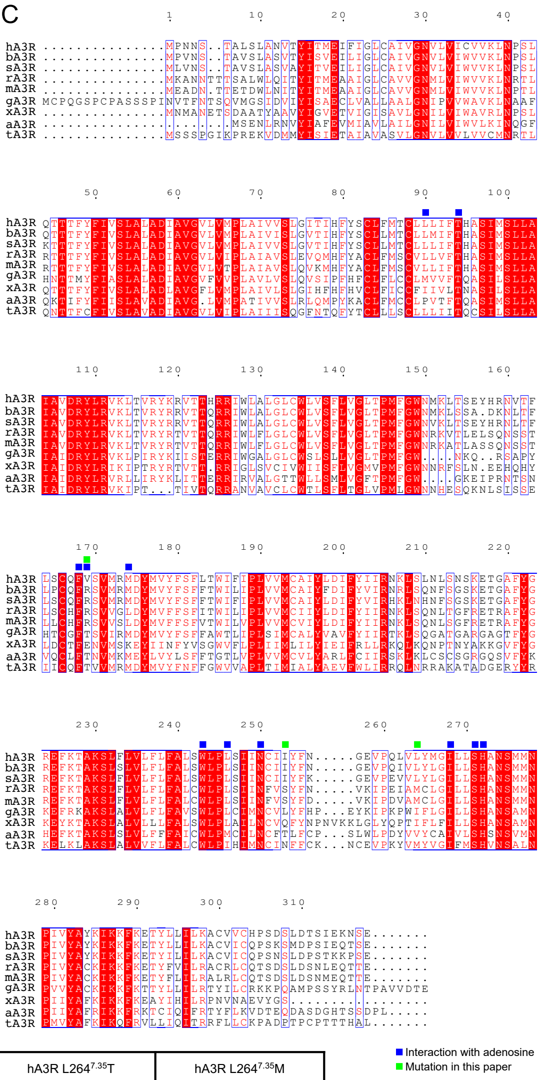


B

| | hA3R WT | | hA3R I253$^{6.58}$T | | hA3R V169$^{5.30}$E | | hA3R L264$^{7.35}$T | | hA3R L264$^{7.35}$M | |
|---|---|---|---|---|---|---|---|---|---|---|
| Ligand | $EC_{50}$ (nM) | Emax (%) | $EC_{50}$ (nM) | Emax (%) | $EC_{50}$ (μM) | Emax (%) | $EC_{50}$ (nM) | Emax (%) | $EC_{50}$ (nM) | Emax (%) |
| m$^6$A | 9.8 ± 0.39 | 43 ± 2.1 | 20 ± 1.8 | 41 ± 2.2 | 2.7 ± 0.15 | 22 ± 2.0 | 6.4 ± 0.44 | 27 ± 2.2 | 28 ± 2.0 | 43 ± 1.7 |
| A | 91 ± 11 | 39 ± 0.82 | 72 ± 2.8 | 37 ± 2.0 | 3.9 ± 0.46 | 14 ± 4.1 | 42 ± 3.3 | 22 ± 0.84 | 92 ± 5.4 | 39 ± 2.6 |

| | hA1R WT | | hA1R T257$^{6.58}$I | | hA1R E172$^{5.30}$V | | hA1R T270$^{7.35}$L | |
|---|---|---|---|---|---|---|---|---|
| Ligand | $EC_{50}$ (nM) | Emax (%) | $EC_{50}$ (nM) | Emax (%) | $EC_{50}$ (nM) | Emax (%) | $EC_{50}$ (nM) | Emax (%) |
| m$^6$A | 840 ± 130 | 17 ± 2.2 | 29 ± 3.6 | 42 ± 1.2 | 110 ± 19 | 32 ± 1.9 | 540 ± 62 | 15 ± 1.6 |
| A | 76 ± 12 | 16 ± 2.5 | 280 ± 36 | 38 ± 1.3 | 110 ± 6.0 | 30 ± 2.9 | 38 ± 6.3 | 11 ± 1.6 |

| | hA2AR WT | | hA2AR T256$^{6.58}$I | | hA2AR E169$^{5.30}$V | | hA2AR M270$^{7.35}$L | |
|---|---|---|---|---|---|---|---|---|
| Ligand | $EC_{50}$ (μM) | Emax (%) | $EC_{50}$ (μM) | Emax (%) | $EC_{50}$ (μM) | Emax (%) | $EC_{50}$ (μM) | Emax (%) |
| m$^6$A | 13 ± 1.1 | 17 ± 0.57 | 3.5 ± 0.63 | 30 ± 1.7 | 4.6 ± 0.88 | 29 ± 2.0 | 1.2 ± 0.56 | 3.5 ± 0.68 |
| A | 1.0 ± 0.063 | 20 ± 0.96 | 13 ± 0.39 | 22 ± 1.1 | 6.4 ± 0.60 | 25 ± 0.76 | N/A | N/A |

D

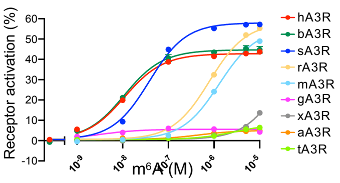


**Figure S5. Molecular basis of selective A3R activation by m$^6$A.**

(A) Amino acid sequence alignment of human A1R (Uniprot ID: P30542), A2AR (Uniprot ID: P29274) and A3R (Uniprot ID: P0DMS8). Secondary structure elements for α-helices and β-strands in the Gi-complexed human A1R are indicated by cylinders and arrows, respectively. Conservation of residues among the three receptors is indicated as follows: red panels for complete conservation; red letters for partial conservation; and black letters for no conservation. The residues involved in adenosine binding are indicated by blue squares, and the residues mutated in this paper are indicated by green squares.

(B) AP-TGFα release responses of adenosine receptor mutants. $EC_{50}$ and $E_{max}$ (%AP-TGFα release) values of adenosine receptor mutants determined by the TGF-α shedding assay. Parameters (means ± SEM) obtained from three independent experiment are shown. The calculated RAi values are shown in Figure 5C.

(C) Alignment of A3Rs across vertebrates. Amino acid sequence alignment of A3R from human (Uniprot ID: P0DMS8), cow (Uniprot ID: Q0VC81), pig (Uniprot ID: F1SBN8), rat (Uniprot ID: P28647), mouse (Uniprot ID: Q3U4C5), chicken (Uniprot ID: R4GIJ8), *Xenopus* (Uniprot ID: ENSXETT00000046736.1), lizard (Uniprot ID: G1KF19), and

*Fugu* (Uniprot ID: H2VBI6). Conservation of the residues among the nine receptors is indicated as follows: red panels for complete conservation; red letters for partial conservation; and black letters for no conservation. Residues involved in adenosine binding are indicated by blue squares, and the residues mutated in this paper are indicated by green squares.

(D) $m^6A$-mediated A3R activation in various species. TGF-α shedding assay of $m^6A$ for A3Rs from the indicated mammals. Initial values were fitted to a four-parameter sigmoidal concentration-response curve using the Prism 8 software. Symbols and error bars are means ± SEM of representative experiments with each performed in triplicate.

Table S1. MRM transition parameters used in this paper

| Metabolites | Precursor ion (m/z) | Product ion (m/z) |
|---|---|---|
| Y | 245.1 | 209 |
| D | 247 | 115 |
| $m^3U$ | 259.1 | 127 |
| U | 245 | 113 |
| C | 244.2 | 112.1 |
| G | 284 | 152.05 |
| $m^3C$, $m^5C$ | 258 | 126 |
| I | 269.1 | 137.1 |
| Cm | 258.25 | 112.05 |
| $m^1G$, $m^2G$, $m^7G$ | 298 | 166 |
| $m^1A$, $m^6A$ | 282.3 | 150.05 |
| $m^1I$ | 283 | 151 |
| Um | 259 | 113 |
| Gm | 298.1 | 152.1 |
| $ac^4C$ | 286.2 | 154 |
| Im | 283.1 | 137.1 |
| $m^2_2G$ | 312 | 180 |
| A | 268 | 136 |
| $t^6A$ | 413.1 | 281.1 |
| Am | 282.1 | 136 |
| $ms^2t^6A$ | 459.1 | 327.1 |
| $m^6_2A$ | 296 | 164 |
| $i^6A$ | 336.2 | 204 |
| $ms^2i^6A$ | 382.2 | 182 |
| $f^5C$ | 272.2 | 140.2 |
| $acp^3U$ | 346.1 | 214 |
| $m^6Am$ | 296 | 150.05 |
| LMS | 182.2 | 56.2 |
